# Local and global chemical shaping of bacterial communities by redox potential

**DOI:** 10.1101/2021.10.12.464155

**Authors:** Jeffrey M. Dick, Delong Meng

## Abstract

Thermodynamics predicts a positive correlation between environmental redox potential and oxidation state of molecules; if found for microbial communities it would imply a new kind of deterministic eco-evolutionary process. This study examines evidence for local- and global-scale correlations between oxidation-reduction potential (ORP or Eh) in environmental samples and carbon oxidation state (*Z*_C_) of estimated bacterial and archaeal community proteomes. Seventy-nine public datasets for seven environment types (river & seawater, lake & pond, alkaline spring, hot spring, groundwater, sediment, and soil) were analyzed. Taxonomic abundances inferred from high-throughput 16S rRNA gene sequences were combined with NCBI Reference Sequence (RefSeq) proteomes to estimate the amino acid compositions and chemical formulas (C_*c*_H_*h*_N_*n*_O_*o*_S_*s*_) of community proteomes, which yield *Z*_C_. Alkaline hot springs have the lowest *Z*_C_ for both bacterial and archaeal domains of any environment. Positive global correlations between redox potential and *Z*_C_ are found for bacterial communities in lake & pond, groundwater, and soil environments, but not archaeal communities, suggesting a broad ecological signal of chemical shaping in Bacteria.

## Introduction

Redox potential, abbreviated as Eh or ORP, is a measure of oxidizing power in a chemical system. Redox potential has been used as a monitor of microcosm development (Pagaling et al., 2014) and ecosystem health (Lanzén et al., 2021) and has important roles for assessing bioremediation (Li et al., 2021; Zecchin et al., 2021) and soil disinfestation (Poret-Peterson et al., 2020) strategies. Although measurements in natural systems generally reflect “mixed potentials” that have no clear thermodynamic interpretation (Bricker, 1982; Tan et al., 2017), the appearance of Eh in numerous microbial ecology studies calls for a better theoretical development of the ecological relevance of this parameter. The primary advantage of Eh for a global-scale assessment of redox effects in microbial communities is that it can be measured over extreme ranges of redox conditions. In contrast, oxygen and hydrogen reach unmeasurable concentrations in anoxic and oxic environments, respectively, which makes either one of them alone less useful as a universal scale of redox conditions.

Redox potential is often one of a number of physicochemical variables that is associated with differences in microbial community composition. Although multivariate techniques can be used to establish correlations between microbial community structure and physicochemical variables, these methods do not have a foundation in chemical concepts, and therefore remain correlative in nature (Fig. 1). Perhaps a hybrid approach that combines elements of biology and chemistry and is informed by the tradition of thermodynamic modeling in geochemistry could shed light on the eco-evolutionary implications of redox potential.

**Fig. 1.**
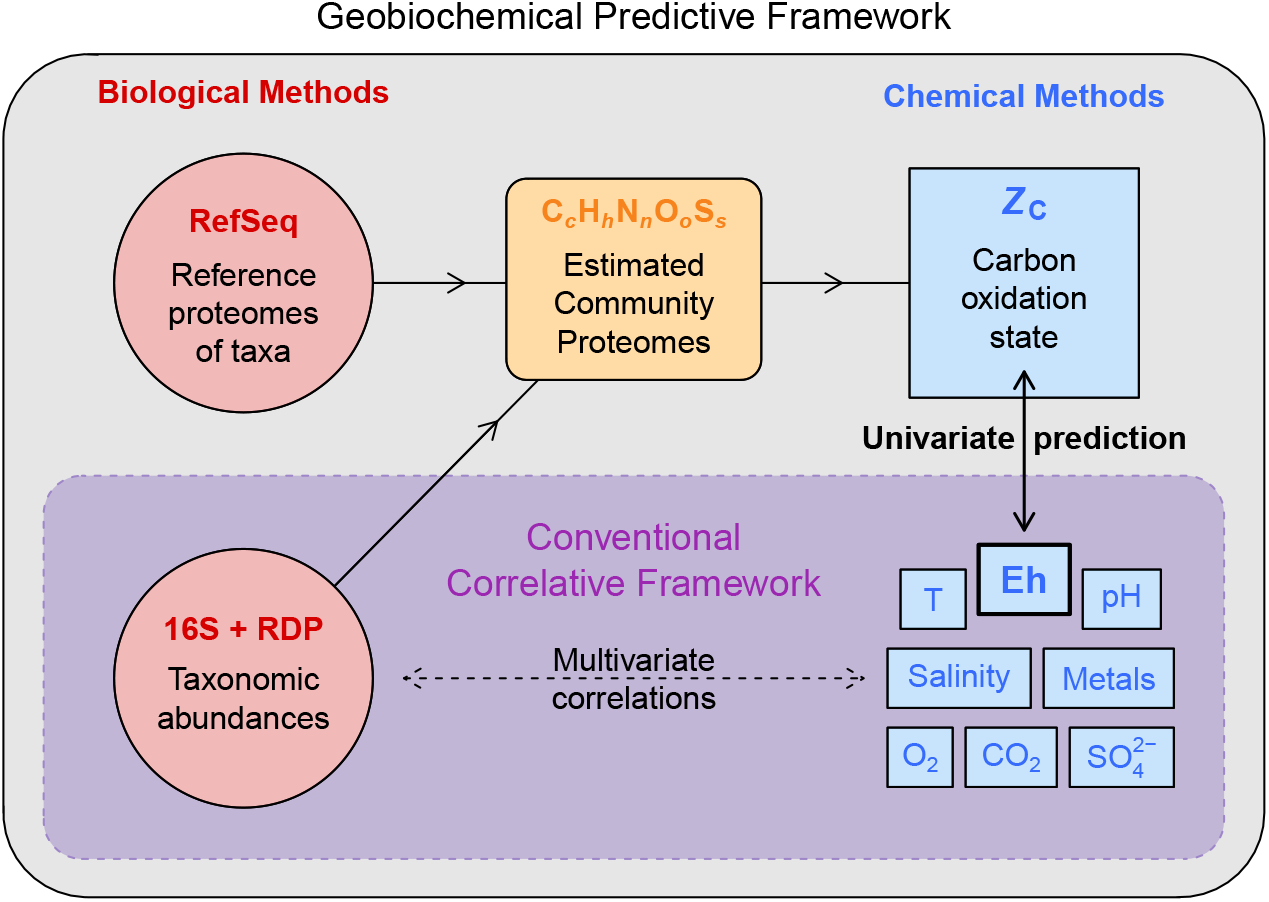
Conceptual framework of the study. This study is concerned with testing the prediction that carbon oxidation state of estimated community proteomes is positively correlated with environmental redox potential. This approach differs from multivariate analyses that identify correlations between taxonomic composition and multiple physicochemical variables but, in general, have no predictive basis.

We propose the term “chemical shaping” for the hypothesis that the elemental composition of microbial communities is linked to environmental redox conditions. Testing this hypothesis requires elemental compositions, which are available for selected microbial species grown in the laboratory (see Popovic et al., 2021), but not for microbial communities in natural environments. To overcome this limitation, it can be recognized that gene sequences carry chemical information in the elemental compositions of the proteins they code for (Dick and Tan, 2021). When microbial abundances inferred from 16S rRNA gene sequences are coupled with reference proteomes of organisms with sequenced genomes, this permits an estimate of the chemical formula of the community proteome, represented as C_*c*_H_*h*_N_*n*_O_*o*_S_*s*_. A basic chemical metric, oxidation state of carbon (*Z*_C_), can be computed for proteins as it can for any organic molecule; for the primary sequence of proteins the equation is (Dick et al., 2020)

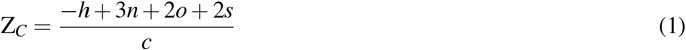

Because *Z*_C_ is a linear combination of elemental ratios, it is a dimensionless quantity, but it can be conceived as a molar electron / carbon ratio, or *e*^-^ / C.

The thermodynamic notion of relative stability allows predicting the direction of irreversible reactions, making it an extremely valuable tool in geochemistry. Completely analogous to how the presence of oxygen increases the relative stability of oxidized minerals compared to more reduced ones, the chemical shaping hypothesis predicts that the redox potential of environments and the oxidation state of biomass are positively correlated. This is specifically not a hypothesis about biochemical reaction mechanisms, but is about differences in (chemical) bulk composition and also about ecology, because the *Z*_C_ of estimated community proteomes is a function of taxonomic composition. To test the prediction, we analyzed data from 79 studies that report both 16S rRNA sequences and redox potential measurements for samples from environment types grouped as River & Seawater, Lake & Pond, Hot Spring, Alkaline Spring, Groundwater, Sediment, and Soil. For estimated proteomes computed from bacterial abundances, we found positive correlations between *Z*_C_ and Eh at three levels: 1) locally within many individual datasets, 2) globally for all datasets for some environment types, and 3) globally for all analyzed datasets for all environments. In contrast to bacterial communities, archaeal communities more often show a negative or no correlation between *Z*_C_ and Eh. Although important variables such as pH, temperature, and salinity are not included in the present analysis, this step toward a chemical representation of microbial communities allows quantifying the physicochemical structuring of communities along redox gradients, uncovers chemical differences between phylogenetic lineages, and may help to build more predictive ecological models.

## Materials and Methods

Analysis of data was performed in three stages: 1) curation of a database for redox potential and sample metadata, 2) sequence data processing and taxonomic classification, and 3) reproducible data analysis and visualization.

### Data sources and Eh corrected to pH 7

In general, datasets were considered for inclusion in the analysis if demultiplexed 16S rRNA sequence data could be found on the Sequence Read Archive (SRA) (Leinonen et al., 2011) or MG-RAST (Meyer et al., 2008) and data were available for at least 5 biological samples with an associated Eh range of at least 100 mV before correction to pH 7. With some exceptions, studies were included where Eh values for individual samples, rather than for means of sample groups, were available. Brief descriptions, BioProject or MG-RAST accession numbers, and references for the datasets are listed in Table 1.

**Table 1.** Datasets analyzed in this study. The name of each dataset is followed by the BioProject or MG-RAST accession number(s) and one or two literature references; in most cases the second reference is for redox potential data. “NA” indicates data that were not obtained from SRA or MG-RAST (see text for details).

| River & Seawater |
| --- |
| Pearl River Estuary (PRJNA319446 and PRJNA357334; Mai et al., 2018), Black Sea (PRJNA423140; Sollai et al., 2019 and van Vliet et al., 2021), Sansha Yongle Blue Hole (PRJNA503500; He et al., 2020 and Xie et al., 2019), Han River Cyanobacterial Blooms (PRJNA530625 and PRJNA594996; Kim et al., 2020), Bahe River (PRJNA588356; Wang et al., 2021), Lancang River (PRJNA636956; Luo et al., 2020), Lagoon of Venice (PRJNA656714; Juhmani et al., 2020), Maozhou River (PRJNA681688; Zhang et al., 2021a), Three Gorges Reservoir (PRJNA733826; Gao et al., 2021) |
| Lake & Pond |
| Kiritimati Microbial Mat (PRJNA174394; Schneider et al., 2013), Siguniang Mountain (PRJNA255556; Li et al., 2017), Andean Lakes (PRJNA289691; Echeverría-Vega et al., 2018), Xidong Reservoir, Xiamen (PRJNA315049; Liu et al., 2019), Potentilla Lake (PRJNA321351; Schütte et al., 2016), Ursu Lake (PRJNA395513; Baricz et al., 2021), Xiamen Reservoirs and Ponds (PRJNA407260; Hu et al., 2018), Mid-Atlantic Agricultural Pond (PRJNA473136; Chopyk et al., 2020), Keweenaw Waterway (PRJNA489447; Butler et al., 2019), Finnish Lakes (PRJNA516525; Rissanen et al., 2021), Fayetteville Green Lake (PRJNA641339; Block et al., 2021), Iberian Pyrite Belt (PRJEB38636; van der Graaf et al., 2020), Lake Kinneret (PRJEB39923; Ninio et al., 2021), Lake Sonachi (PRJNA731062; Fazi et al., 2021), Peña de Hierro (mgp83146; Grettenberger et al., 2020) |
| Hot Spring |
| Bison Pool, Yellowstone National Park (NA; Swingley et al., 2012 and Dick and Shock, 2011), New Zealand Geothermal Springs (PRJEB24353; Power et al., 2018), Dallol Geothermal Area (PRJNA541281; Belilla et al., 2019), Copahue Volcano-Río Agrio (PRJNA542136; Lopez Bedogni et al., 2020 and Massello et al., 2020), Southwestern Yunnan (PRJNA548851; Guo et al., 2021), Eastern Tibetan Plateau (PRJNA592622; Guo et al., 2020), Uzon Caldera (PRJNA623081; Peltek et al., 2020), Southern Tibetan Plateau (PRJNA638734; Ma et al., 2021), Japan Hydrothermal Sediment (NA, Omae et al., 2019) |
| Alkaline Spring |
| Coast Range Ophiolite Microbial Observatory (CROMO) (PRJNA289273; Sabuda et al., 2020), Samail Ophiolite (PRJNA352492; Rempfert et al., 2017), Santa Elena Ophiolite (PRJNA361138; Crespo-Medina et al., 2017), Voltri Massif (PRJNA685937; Kamran et al., 2021), Samail Ophiolite Packers (PRJNA743134; Nothaft et al., 2021) |
| Groundwater |
| Sarnia nZVI Injection (PRJNA308958; Kocur et al., 2016), Mahomet Aquifer (PRJDB5959; Yamamoto et al., 2019), Pearl River Delta (PRJNA408058; Sang et al., 2019), Rayong Province (PRJNA434769; Sonthiphand et al., 2019), Mezquital Valley (PRJNA488796; Aguilar-Rangel et al., 2020); Phetchabun-Pichit Gold Mine (PRJNA528471; Sonthiphand et al., 2021a), Hainich Critical Zone (PRJEB33032; Yan et al., 2020), Zoovch Ovoo Uranium Deposit (PRJNA553521; Jroundi et al., 2020), Golmud, Qaidam Basin (PRJNA623081; Guo et al., 2019), Lower Chao Praya Basin (PRJNA630252; Sonthiphand et al., 2021b), Po Plain (PRJNA667833; Zecchin et al., 2021) |
| Sediment |
| Loki's Castle (SRP009131; Jorgensen et al., 2012), Humboldt Sulfuretum (PRJNA262691; Gallardo et al., 2016), Mai Po Wetland (PRJEB12432 and PRJEB12429; Zhou et al., 2017), Great Salt Lake (PRJNA319444; Boyd et al., 2017), Baltic Sea Swedish Coast (PRJNA322450; Broman et al., 2017), Honghu Lake (PRJNA352457; Han et al., 2019), Daya Bay (PRJNA400089; Wu et al., 2021), Perdido Fold Belt (PRJNA429278; Sánchez-Soto Jiménez et al., 2018), Jinchuan River (PRJNA437688, PRJNA437695, PRJNA437692, and PRJNA437697; Cai et al., 2019), Loa River (PRJEB28365; Zárate et al., 2020), Cock Soda Lake (PRJNA453733; Vavourakis et al., 2019), Qiangtang River Estuary (PRJNA488529; Wang et al., 2019), Lake Neusiedl (PRJNA507590; von Hoyningen-Huene et al., 2019), Valparaíso Bay (PRJEB31703; Molina et al., 2021), Bay of Biscay (PRJEB35647; Lanzén et al., 2021), Maozhou River Hyporheic Zone (PRJNA616197; Zhang et al., 2021b), Estuarine Sediment Mesocosms (PRJNA639965; Wyness et al., 2021) |
| Soil |
| Biochar in Paddy Soil (PRJNA341915; Song et al., 2017), Dabu Town, Hunan Province (PRJNA361046; Meng et al., 2019), Urban Wetland Soils (PRJNA415514; Brigham et al., 2018), Pentachlorophenol Polluted Soil (PRJNA427749 and PRJNA427853; Zhu et al., 2018), Soil Drainage and Imbibition (PRJEB28313; Pronk et al., 2020), Hydrocarbon Biodegradation (PRJNA523725; Zhang et al., 2019), Paddy Soil Water Management (PRJNA564714; Chuang et al., 2020), Anaerobic Soil Disinfestation (PRJNA575041; Poret-Peterson et al., 2020), Cd Bioleaching Remediation (PRJNA576993; Xu et al., 2020), East Asia Paddy Soil (PRJNA607877; Luan et al., 2020), Rice Rhizosphere (PRJNA611687; Dai et al., 2020), Reductive Soil Disinfestation (PRJNA665924; Li et al., 2021), Paddy Soil Amendments (PRJNA690162; Dykes et al., 2021 and Leinonen et al., 2011) |

Values of redox potential, assumed to be stated with respect to the standard hydrogen electrode (SHE), were taken from the original publications as detailed in Table S1. Where needed, values were extracted from plots using the program g3data (Frantz and Novak, 2013). Values of temperature and pH were also compiled if available; otherwise their values were assumed to be 25 °C and pH 7 for the conversion to Eh7. Occasionally, redox potential values were obtained from different publications than those that describe the sequencing data; see Table 1 for references and Table S1 for additional details. Textual descriptions of sample type (e.g. water or sediment), location (e.g. geographic location or depth in water or sediment columns), sampling time (e.g. date or season), or treatment group for soil and sediment mesocosms were also recorded as applicable to each study.

Because of the dependence of electrical potential on pH – known as the Nernst slope or Nernst factor – reported Eh values were corrected to values at pH 7 with the equation (Husson et al., 2016)

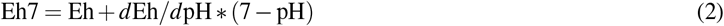

where Eh7 denotes the corrected value and *d*Eh*/d*pH is the theoretical slope, which is equal to −59.16 mV/pH unit at 25 °C. Values of the Nernst slope at other temperatures were calculated from −2.303*RT*/*F*, where *R, T*, and *F* are the gas constant, temperature in Kelvin, and Faraday constant.

### Sequence data processing and chemical metrics

In most cases, one sequencing run per sample was used. The biological material was DNA except for the Great Salt Lake sediment study (Boyd et al., 2017), which used RNA reverse transcribed to cDNA. For Bison Pool Hot Spring in Yellowstone National Park (Swingley et al., 2012), 16S rRNA sequence data were obtained from the authors of that study. The dataset of Power et al. (2018) includes 16S rRNA sequence data for 925 water samples of hot springs in New Zealand; for this study we used data for randomly selected 42 acidic samples (pH *<* 3) and 39 circumneutral to alkaline samples (pH > 6). A multi-year dataset with 256 samples is available for sediments in the Bay of Biscay (Lanzén et al., 2021); for this study, we used data for 55 samples collected in 2017.

Sequence data were processed with a pipeline consisting of removal of adapter sequences, merging of forward and reverse reads where applicable, quality and length filtering, singleton removal, subsampling to 10000 sequences for each sample, reference-based chimera detection, and taxonomic classification. Length filtering was specified as a minimum read length of 200 and maximum length of 600 for 454 pyrosequencing datasets (--fastq minlen and --fastq maxlen options of vsearch), or as a truncation length for Illumina datasets (--fastq trunclen option), which was manually selected for each dataset as approximately the highest round-number value that could be used while retaining the majority of reads. Specific settings used for each dataset are given in Table S2. After length, quality, and singleton filtering, runs were subsampled at a depth of 10000 sequences so that the chimera detection and taxonomic classification steps could be performed in a reasonable amount of time. The pipeline depends on the following software and databases: fastq-dump from the NCBI SRA Toolkit with the --clip option to remove adapter sequences (has effect only for 454 datasets), vsearch (Rognes et al., 2016) for merging, singleton and chimera detection, seqtk (Li, 2020) for extracting non-singletons, SILVA SSURef NR99 database version 138.1 (Quast et al., 2012) for chimera detection, and RDP Classifier version 2.13 (Cole et al., 2014) for taxonomic classification with the default confidence threshold of 80%. After classification using the -h option of the RDP Classifier to produce counts of reads assigned to each taxon, the results for all samples in a dataset were merged using the merge-count command of the RDP Classifier.

The -h option of the RDP Classifier produces a hierarchical file where sequences are assigned at multiple taxonomic levels. For subsequent processing, assignments at the root or domain level or to Chloroplast or Eukaryota were omitted, and the lowest assigned taxonomic level (from genus to phylum) for each remaining sequence was used for mapping. Taxonomic classifications in the RDP output were mapped to the NCBI taxonomy represented in RefSeq by automatic name matching or by custom mappings for particular taxa; see Dick and Tan (2021) for details. Unmapped taxonomic classification were omitted from subsequent analysis. Reference proteomes of archaeal and bacterial taxa at levels from genus to phylum in the NCBI Reference Sequence (RefSeq) database release 206 (O’Leary et al., 2016) generated by combining amino acid compositions for all proteins in each taxon were described previously (Dick and Tan, 2021) and used here. Amino acid compositions were used to calculate *Z*_C_ for each taxon accounting for different carbon numbers of amino acids (Dick et al., 2020).

For one dataset for hydrothermal sediment in Japan (Omae et al., 2019), sequence data processing was not done in this study. Instead, the table of 13798 OTUs in Online Resource S2 of Omae et al. (2019) was filtered to keep the 709 most abundant OTUs (those with read counts *>* 500 across all samples) and converted to .tab format similar to the output file of the RDP Classifier merge-count command. The lowest-level taxonomic name for each OTU was assigned a taxonomic rank and was used for mapping to RefSeq; if no mapping was available, the next lowest-level taxonomic name in the OTU table was used; all remaining unmapped OTUs were discarded from subsequent analysis.

To calculate *Z*_C_ values, taxonomic abundances in bacterial and archaeal domains were considered separately. For sequencing runs for libraries generated using domain-specific bacterial or archaeal primers, custom R functions were used to retain only the bacterial or archaeal taxonomic assignments, respectively, in the output of the RDP Classifier. The use of domain-specific primers is indicated by the “Domain” column in the metadata files generated here. If studies used universal primers (often 515F/806R), R functions were used to extract the RDP assignments within each domain. Samples with less than 50 sequences with lowest-level taxonomic assignments from genus-to phylum-level for either Bacteria or Archaea were discarded from further analysis for that domain. According to these criteria, all datasets but one (Wang et al., 2019) have bacterial sequences, and many datasets have both bacterial and archaeal sequences (see Table S1). Within each domain, the abundances of all mappable lowest-level taxonomic assignments were multiplied by *Z*_C_ of the reference proteomes, summed, and divided by total abundance to estimate *Z*_C_ of the bacterial or archaeal community.

### Computer code for data analysis and visualization

The code and output files for the steps described above are available in the JMDplots package on GitHub (https://github.com/jedick/JMDplots). The sequence processing pipeline is implemented in the script `chem16S/pipeline.R`. Metadata including accession numbers, sample ID and description, Eh, *T* and/or pH are saved in .csv files for each study in the `orp16S/metadata` directory. The .tab files produced by the RDP merge-count command are saved in the `orp16S/RDP` directory. The amino acid compositions of reference proteomes, their taxonomic names and ranks, and scripts used to generate these files from the the RefSeq database are in the `refseq` directory (see Dick and Tan, 2021 for details). The `getmdat()` function retrieves metadata for a dataset using a bibliographic key and implements the Eh7 correction, the `getRDP()` function implements the selection of assignment counts at lowest taxonomic levels and includes a “lineage” argument to select e.g. only Bacteria or Archaea, the `getmap()` function maps RDP taxonomic names to NCBI names, and the `getmetrics()` function implements the calculation of *Z*_C_ for the estimated community proteomes. Additional code used for the figures includes R’s `boxplot()` (R Core Team, 2021) for the *Z*_C_ distribution at each depth of Winogradsky columns in Fig. 2B and the oce package (Kelley and Richards, 2021) for the world map in Fig. 3.

**Fig. 2.**
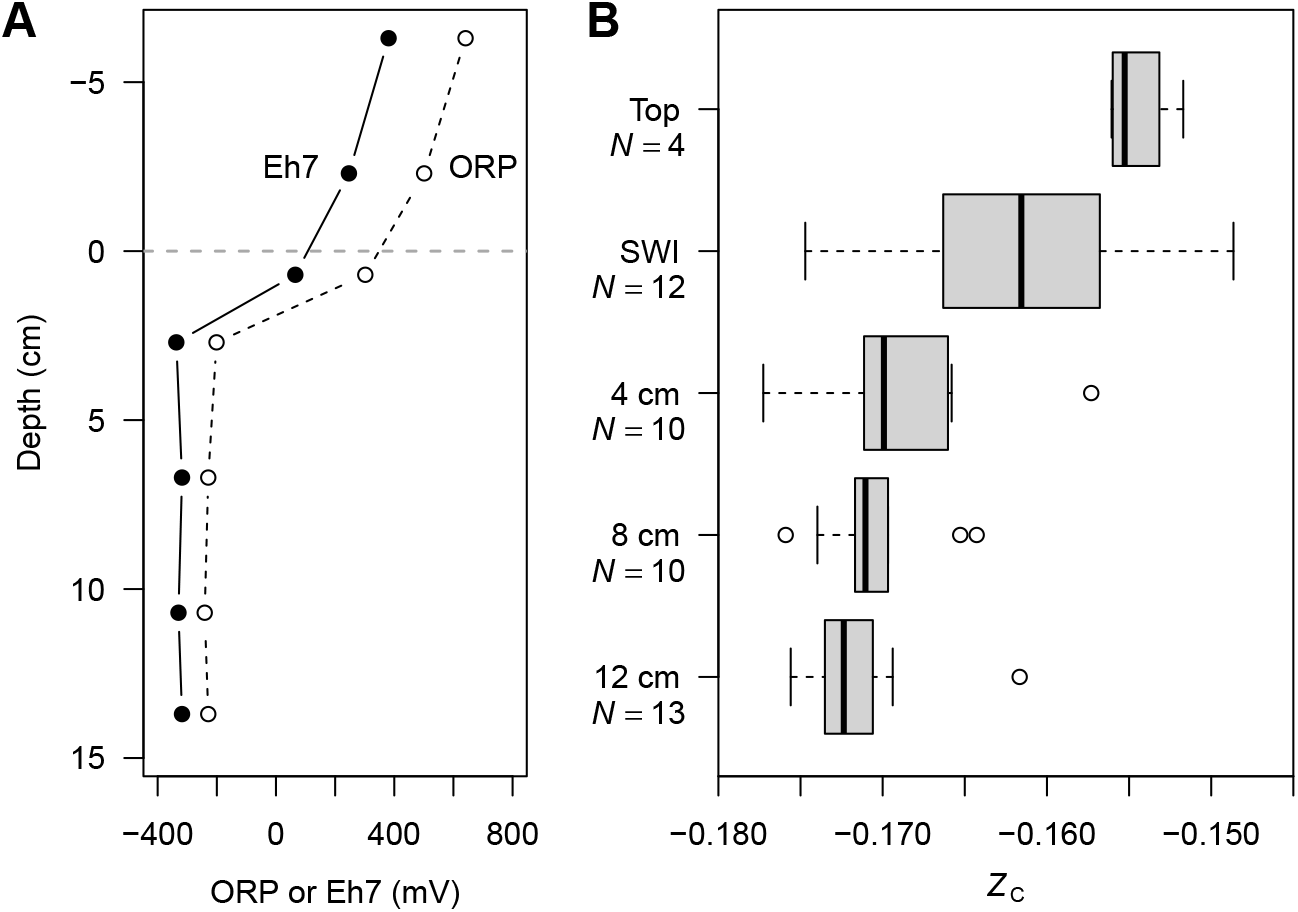
Chemical and geobiochemical depth profiles in Winogradsky columns. (A) ORP measurements (dashed line) in a Winogradsky column, taken from Figure S5 of Diez-Ercilla et al. (2019) with the depth scale adjusted so the sediment-water interface is at 0 cm. Values of Eh7 computed from Eq. (2) are also shown (solid line); they are shifted to more negative values because of the acidic pH in the experiments of Diez-Ercilla et al. (2019). This is a representative situation chosen for the availability of both ORP and pH data at multiple depths in water and sediment; Winogradsky columns made with non-acidic sediment also show decreasing redox potential with depth (see text for references). (B) Values of *Z*_C_ for estimated community proteomes computed using the 16S rRNA data of Rundell et al. (2014) for non-acidic sediment from ponds in Massachusetts, USA (BioProject PRJNA234104). The box-and-whisker plots represent data for the same depth intervals in different columns. “Top” indicates samples collected by scraping the top layer of sediment, while “SWI” denotes samples of the sediment-water interface collected by drilling into the side of the column (Rundell et al., 2014).

**Fig. 3.**
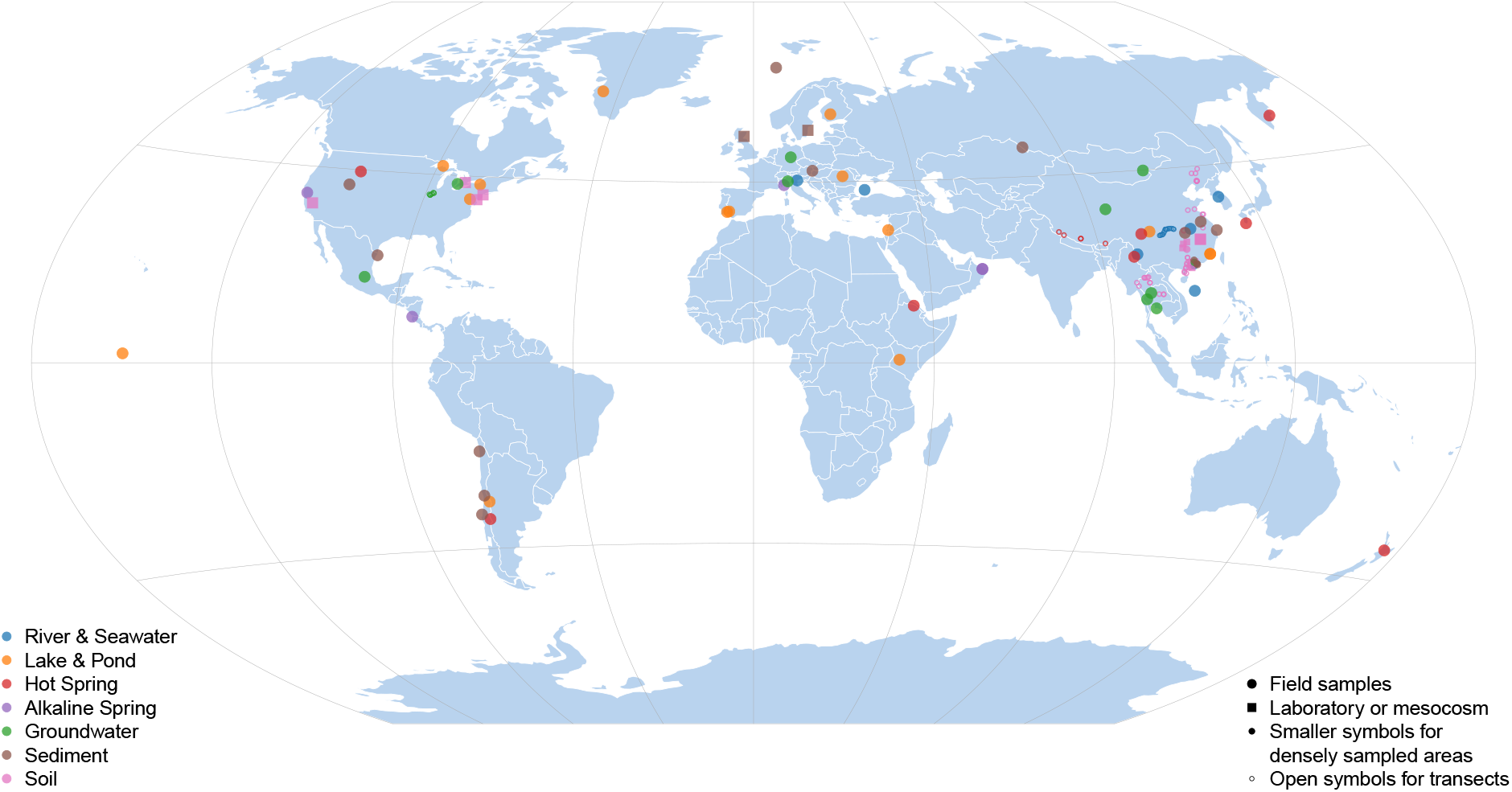
Approximate sampling locations on world map. Colors denote environment types, smaller symbols are used for areas with high densities of sites (Hunan Province in Central South China and the Guangdong–Hong Kong–Macau Greater Bay Area in South China), and small open circles indicate sampling transects for paddy soils in East Asia (Luan et al., 2020), hot springs in the Southern Tibetan Plateau (Ma et al., 2021), water samples from the Three Gorges Reservoir (Gao et al., 2021), and wells in the Mahomet Aquifer, Illinois, USA (Yamamoto et al., 2019). Studies used field-collected samples except for those indicated by square symbols, which are for laboratory experiments or mesocosms. The map is drawn with the Winkel Tripel projection and was created using the R package oce and its included coastlineWorld dataset, together with shapefiles for the North American Great Lakes obtained from USGS (2010). Geographic coordinates of sampling sites were taken from publications or from the BioSample metadata on NCBI; see the source code of the `orp16S3()` function in the JMDplots package for details.

The function `orp16S_S1()` in the JMDplots package is used to make the plots for each dataset in Figure S1 and outputs .csv files for sample data and linear regression coefficients that correspond to Datasets S1 and S2. These datasets are used by other functions in the package to make each figure (e.g. `orp16S4()` for Fig. 4), and the vignette `orp16S.Rmd` runs each function to reproduce the figures in this paper. Studies are identified in the package, supplementary material, and plot titles of Fig. 4 by bibliographic reference keys, which are defined in the BibTeX file in Dataset S3.

**Fig. 4.**
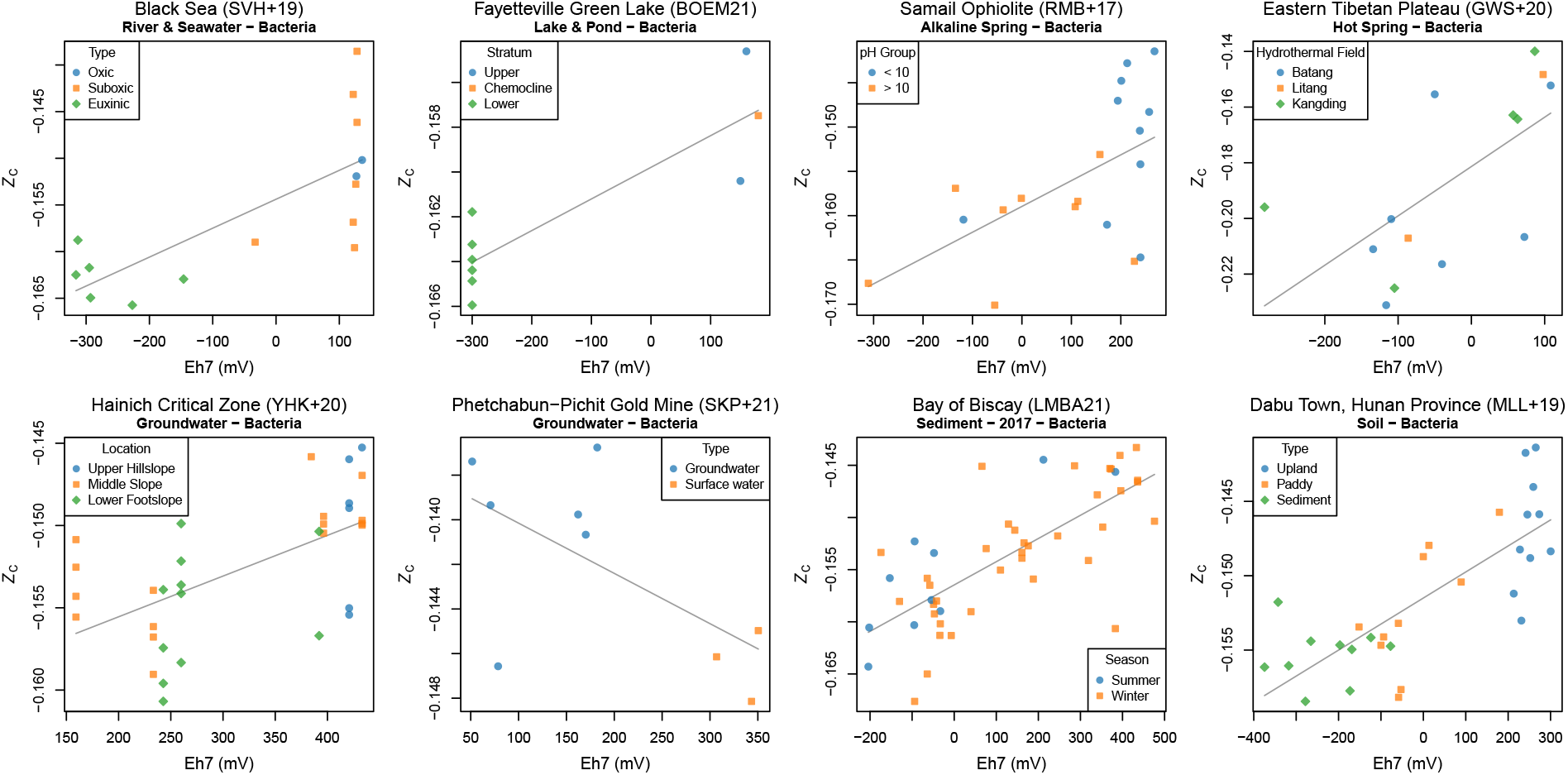
Analysis of selected datasets for each environment type. Plots show sample data and linear regressions between Eh7 and *Z*_C_ for bacterial abundances. Solid lines indicate that the absolute value of the fitted slope is *>* 0.01 V^-1^. See Figure S1 for plots for all datasets.

## Results

### Winogradsky columns

To establish the feasibility of these methods, we analyzed data for Winogradsky columns, which are well known as teaching and research tools to illuminate basic concepts of microbiology by monitoring the biological and chemical changes in enriched sediments and overlying water in a controlled laboratory setting (Rundell et al., 2014). Winogradsky columns are also candidates for “model systems” for microbial communities that may facilitate transferability of results between studies (Widder et al., 2016). As far as we are aware, there are no published studies that provide 16S rRNA data and ORP and pH measurements along depth profiles in the same Winogradsky columns, so we have used data from different sources with the aim of uncovering general features. Therefore, the following analysis is based on the *direction of change* of redox potential and estimated community *Z*_C_ with depth but not their specific values, which likely differ depending on source material and laboratory parameters.

Fig. 2A shows ORP measurements taken from Diez-Ercilla et al. (2019) for a column consisting of acidic sediment and water from pit lakes in the Iberian Pyrite Belt, Spain. Because of the acidic conditions in these experiments, Eh values corrected to pH 7, denoted as Eh7, are shifted to lower values than the reported redox potential, but still show a decreasing trend with depth. Mature Winogradsky columns prepared using sediment from non-acidic ponds also exhibit decreases of redox potential with depth in the sediment (Pibernat and Abellà, 1993; Pagaling et al., 2014), and more reducing conditions in sediment compared to overlying water (see Figure 5 of Widder et al., 2016 and Figure S1A of Pagaling et al., 2017). It follows that a decrease of redox potential with depth is a general feature of mature Winogradsky columns.

**Fig. 5.**
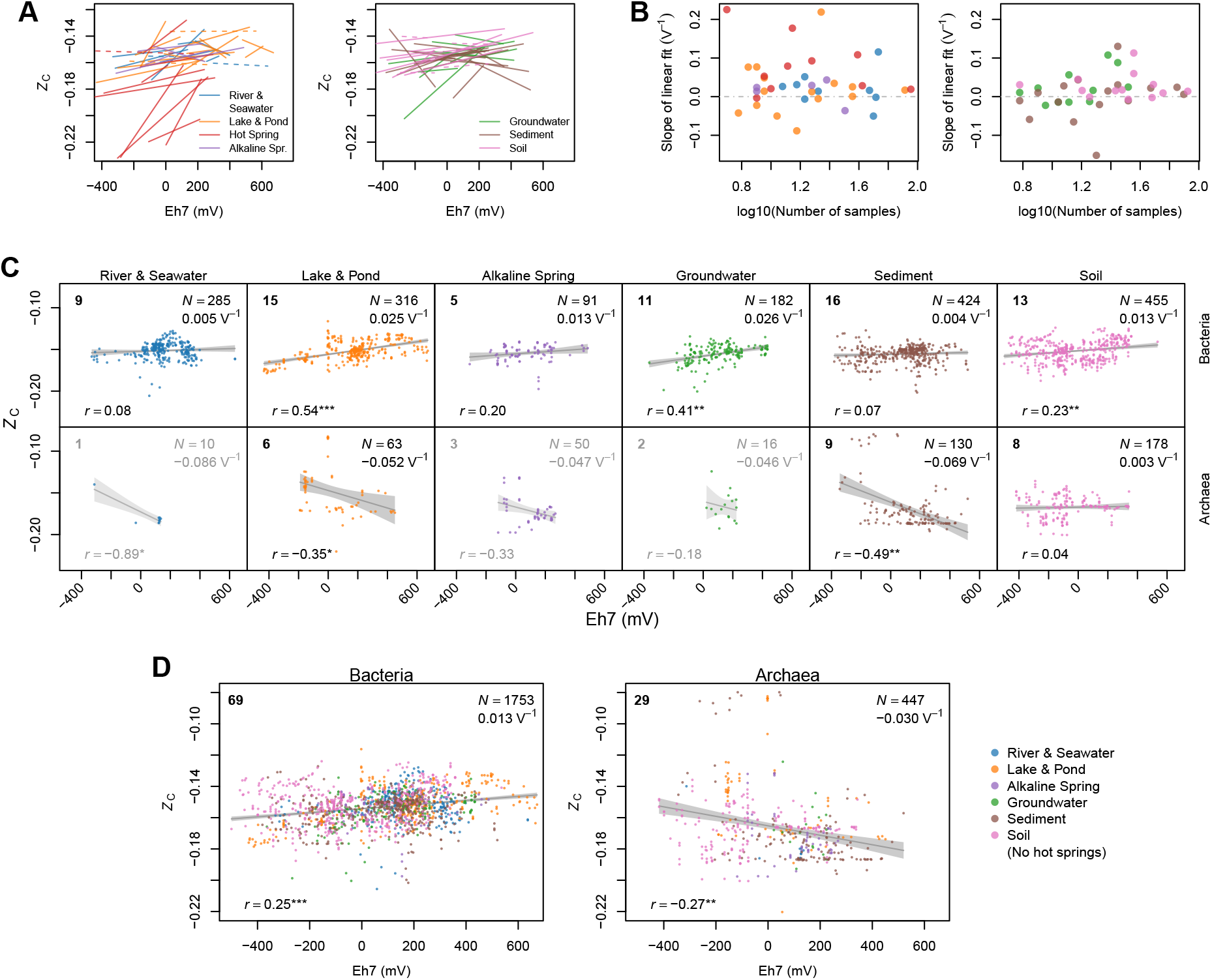
Local and global analysis of *Z*_C_ and Eh7 data. (A) Linear regressions between Eh7 and *Z*_C_ of estimated bacterial community proteomes in each dataset. (B) Slopes of the linear regressions in (A) plotted against the decimal logarithm of number of samples for each dataset. (C) *Z*_C_–Eh7 plots and linear regressions for all data in each environment type. The bold numbers in the upper-left corner give the number of datasets, and the legends in the upper-right corner display the total number of samples (*N*) and slope of the linear regression. Slopes are computed for *Z*_C_ as a function of Eh7 (V), which is 1000 times the slope against Eh7 (mV). The lower-left legends show the Pearson correlation coefficient (*r*) with *P*-value indicated by stars (\**P <* 10^−2^, \*\**P <* 10^−5^, \*\*\**P <* 10^−20^). The text is grayed out for environments with fewer than 5 datasets. (D) *Z*_C_–Eh7 plots and linear regressions for Bacteria and Archaea for all environment types except hot springs.

According to the chemical shaping hypothesis, the elemental composition of microbial communities is predicted to become more reduced with depth in mature Winogradsky columns. Data supporting this prediction are shown in Fig. 2B for *Z*_C_ values calculated using 16S rRNA sequence data from Rundell et al. (2014) for columns made with sediment from ponds in Massachusetts, USA. Rundell et al. (2014) noted characteristic changes in community composition with depth; relatively high abundances of Alphaproteobacteria and Betaproteobacteria occur near the top of the columns, and more abundant Firmicutes, particularly Clostridia, are found in the lower parts. These taxonomic differences are the immediate cause for the decreasing trend of *Z*_C_. Proteobacteria and Clostridia are lineages with relatively oxidized and reduced reference proteomes, respectively (Dick and Tan, 2021); accordingly the estimated *Z*_C_ decreases with depth in the columns. It can also be noted by observing the relative lengths of the whiskers in Fig. 2B that the sediment-water interface (SWI) is the layer with the greatest variability of *Z*_C_. This is concordant with the relatively low similarity of communities in replicate samples for the SWI as revealed by UNIFRAC distances (Rundell et al., 2014) and further emphasizes that *Z*_C_ is a chemical representation of the taxonomic composition of communities.

The decrease of estimated carbon oxidation state with depth in Winogradsky columns supports the thermodynamic prediction that the communities are chemically shaped by redox potential. The rest of this study is devoted to analyzing communities in a wide variety of environments in order to assess the broader applicability and limitations of this prediction.

### Analysis of local-scale data for different environment types

We searched the literature for studies that report both 16S rRNA and redox potential (ORP or Eh) data. Several types of laboratory-based studies were excluded from this search, so that most of the included studies are field-based, but mesocosms and laboratory experiments on soils and some sediments were included.

The locations of study sites are shown in Fig. 3. For each study, a plot of Eh7 and bacterial *Z*_C_ was made; if archaeal sequences were available, a second plot for archaeal *Z*_C_ was made. Figure S1 has scatterplots with linear regressions for all datasets, and selected datasets are shown in Fig. 4. In these figures, solid and dashed lines indicate linear fits with an absolute value of slope greater than or less than 0.01 V^-1^, respectively. In the following paragraphs, datasets are referred to as having a positive (slope *>* 0.01 V^-1^) or negative (slope *<* −0.01 V^-1^) correlation, or no correlation (|slope| *<* 0.01 V^-1^). The linear regressions do not capture complex patterns in the data, but the sign of the slope enables straightforward testing of the prediction that the correlation between Eh7 and *Z*_C_ is positive.

The following paragraphs describe the results in more detail for bacterial communities. The plots in Figure S1 reveal that many archaeal communities show different trends, but for conciseness, the data for Archaea will only be described in the section on global-scale analysis.

#### River & Seawater and Lake & Pond

In multiple datasets for water bodies, spatial variation in redox potential is associated with corresponding changes in *Z*_C_ of the bacterial communities. Several stratified water columns, including the PHOX2 station in the western Black Sea (Sollai et al., 2019), the Sansha Yongle Blue Hole in the South China Sea (He et al., 2020), and Ursu Lake in Romania (Baricz et al., 2021) were previously found to exhibit decreasing *Z*_C_ of estimated community proteomes with depth (Dick and Tan, 2021). In the present study, we find that *Z*_C_ for bacterial communities exhibits a positive correlation with Eh7 calculated from redox potential measurements taken from van Vliet et al. (2021), Xie et al. (2019), and Baricz et al. (2021), respectively (see Figure S1 for plots for all datasets and Fig. 4 for the Black Sea). Other lakes where *Z*_C_ is aligned with the vertical redox potential gradient are Fayetteville Green Lake in New York, USA (Block et al., 2021) (Fig. 4), Lake Sonachi in Kenya (Fazi et al., 2021), Potentilla Lake in Greenland (Schütte et al., 2016), and two lakes in the Iberian Pyrite Belt (IPB) (van der Graaf et al., 2020).

One sample included in the group of lake datasets is not for water, but for the ca. 10 cm-thick microbial mat in a hypersaline lake of the Kiritimati Atoll in the central Pacific Ocean (Schneider et al., 2013). There is a large separation between relatively high *Z*_C_ in the photic-oxic zone near the top of the mat (Eh7 *≈* 0 mV) and lower *Z*_C_ in the transition and anoxic zones toward the bottom (Eh7 *≈* 150 mV).

Geographically separated stations on rivers including the Pearl River Estuary in south China (Mai et al., 2018) and Lancang River in southwestern China (Luo et al., 2020) also exhibit positive slopes for linear fits between *Z*_C_ of the bacterial communities and redox potential. In contrast, negative correlations between Eh7 and bacterial *Z*_C_ are apparent for depth profiles in two Finnish lakes (Rissanen et al., 2021) and Peña de Hierro in the IPB (Grettenberger et al., 2020), temporal variations in an agricultural pond in the Mid-Atlantic region of the USA (Chopyk et al., 2020), water samples along a transect in the Three Gorges Reservoir (Gao et al., 2021), and an elevation gradient in Siguniang Mountain in western China (Li et al., 2017). The negative correlation at Peña de Hierro speculatively could be associated with the unusual geochemistry of this lake compared to others in the IPB, particularly microoxic rather than anoxic conditions at depth (Grettenberger et al., 2020). The other datasets that were analyzed but not mentioned here, including several for time series during cyanobacterial bloom or ice formation, yield linear regressions with an absolute value of slope that is less than 0.01 V^-1^.

#### Alkaline Spring and Groundwater

The Alkaline Spring datasets represent water samples from serpentinizing systems associated with weathering of ultramafic peridotites. Positive correlations between Eh7 and *Z*_C_ are exhibited by samples from the Voltri Massif, Italy (Kamran et al., 2021), which include two spring sites with different ranges of redox potential and a more oxidizing creek sample, samples from wells with different borehole depths in the Coast Range Ophiolite Microbial Observatory (CROMO), California, USA (Sabuda et al., 2020) and the Samail Ophiolite, Oman (Rempfert et al., 2017) (Fig. 4), and depth-resolved water samples from the Samail Ophiolite collected using a packer system (Nothaft et al., 2021). In contrast, there is relatively little variation of bacterial community *Z*_C_ over an Eh7 range of ca. −50 to 250 mV for spring, well, and river samples from the Santa Elena Ophiolite, Costa Rica (Crespo-Medina et al., 2017), except for samples from a single well at relatively high redox potential and low *Z*_C_, which results in an overall negative correlation.

For Groundwater datasets (those not associated with serpentinization), positive correlations between bacterial *Z*_C_ and Eh7 are found for samples from roll-front redox zones of the Zoovch Ovoo Uranium Deposit (Jroundi et al., 2020), a time series following injection of nanoscale zerovalent iron (nZVI) in Sarnia, Ontario, Canada (Kocur et al., 2016), wells in areas with mixed land uses in Rayong Province, Thailand (Sonthiphand et al., 2019), a hillslope transect in the Hainich Critical Zone, Germany (Yan et al., 2020), wells along a groundwater salinity gradient in the Pearl River Delta, China (Sang et al., 2019), wells in the Mahomet Aquifer, Midwestern USA (Yamamoto et al., 2019), and samples from a canal, piezometer, well, and spring in Mezquital Valley, Mexico (Aguilar-Rangel et al., 2020). In the latter dataset, the canal has the lowest range of both measured redox potential and estimated bacterial *Z*_C_, which extends to lower values than the other datasets for groundwater and other environment types, except those for alkaline hot springs (Fig. 5A).

In contrast, there is a negative correlation between bacterial *Z*_C_ and Eh7 for groundwater and surface water samples from the Golmud area in the Qaidam Basin, China (Guo et al., 2019), in the vicinity of a gold mine in Phetchabun and Pichit provinces in Thailand (Sonthiphand et al., 2021a) (Fig. 4), among shallow and deep groundwater samples in Lower Chao Praya Basin (Sonthiphand et al., 2021b), and in the Po Plain, Northern Italy (Zecchin et al., 2021). Notably, arsenic was identified as a major driver of community structure in several of these studies (Sonthiphand et al., 2021a, 2019; Zecchin et al., 2021). A novel hypothesis to emerge from the present analysis is that arsenic contamination may interfere with the redox-based chemical shaping of the bacterial communities.

#### Sediment and Soil

The 15 field-based studies of sediments are approximately evenly divided between those with positive and negative correlations. The datasets with positive correlations include some for marine (Jorgensen et al., 2012; Gallardo et al., 2016), coastal/estuarine (Lanzén et al., 2021) (Fig. 4), freshwater (Han et al., 2019; Cai et al., 2019; Zhang et al., 2021b), and hypersaline (Boyd et al., 2017) sediments. Several of these reflect depth profiles in sediment cores (Jorgensen et al., 2012; Gallardo et al., 2016; Zhou et al., 2017), whereas others include both sediment and water samples (Han et al., 2019), and one study investigated depth profiles in both the water column and sediment of the Jinchuan River in China (Cai et al., 2019).

The datasets with negative correlations represent marine (Sánchez-Soto Jiménez et al., 2018; Molina et al., 2021; Wu et al., 2021), coastal (Zhou et al., 2017), freshwater (Zárate et al., 2020; von Hoyningen-Huene et al., 2019), and hypersaline sediments (Vavourakis et al., 2019). The dataset with the most negative fitted slope is that for Daya Bay, China (Wu et al., 2021), where samples were collected from the surface, middle, and bottom of sediment cores. In addition, multiple lake and ocean datasets are characterized by higher Eh7 for overlying water than sediment (Han et al., 2019; von Hoyningen-Huene et al., 2019; Vavourakis et al., 2019; Molina et al., 2021); most of these exhibit negative correlations with *Z*_C_, except for Honghu Lake (Han et al., 2019), which shows a relatively high *Z*_C_ for water. A systematic difference in redox potential between water and sediment is not evident for river datasets (Cai et al., 2019; Zárate et al., 2020).

Two datasets from laboratory experiments with sediments show positive correlations between Eh7 and *Z*_C_: sediments and water from the Baltic Sea Swedish Coast subjected to incubation under oxic or anoxic conditions (Broman et al., 2017) and depth profiles in estuarine sediment mesocosms with different macroinvertebrate ecosystem engineers (Wyness et al., 2021).

Most of the soil datasets considered here (11 of 13) were obtained from laboratory or mesocosm studies. The absolute values of the *Z*_C_–Eh7 slope for two of these are less than 0.01 V^-1^ (Zhu et al., 2018; Xu et al., 2020), and the remainder show positive correlations between Eh7 and *Z*_C_. The experimental designs include biochar addition in soil remediation experiments (Song et al., 2017), acetate addition to urban wetland soil (Brigham et al., 2018), water management and cyclic drainage and imbibition (Pronk et al., 2020; Chuang et al., 2020), anaerobic or reductive soil disinfestation (Poret-Peterson et al., 2020; Li et al., 2021), inoculation with soil of an artificially petroleum hydrocarbon-contaminated sediment with different terminal electron acceptors (Zhang et al., 2019), sampling of bulk soil and mature, elongation, and tip zones of rhizosphere of rice plants (Dai et al., 2020), and amendment of paddy soils with rice husk, charred husk, or calcium silicate (Dykes et al., 2021).

There are two field-based soil datasets. East Asia Paddy Soil sampled along a regional transect (Luan et al., 2020) has a slightly negative fitted slope between Eh7 and *Z*_C_. A large positive correlation is evident for samples of Cd-contaminated upland soil, paddy soil, and sediment in Dabu Town, Hunan Province, China (Meng et al., 2019) (Fig. 4).

### Global-scale analysis

Linear regressions between Eh7 and bacterial *Z*_C_ for all datasets are compiled in Fig. 5A in separate panels for (1) River & Seawater, Lake & Pond, Alkaline Spring, Hot Spring and (2) Groundwater, Sediment, Soil. The slopes are positive for most datasets in the first group and also for most groundwater and soil datasets. This can be seen more easily in Fig. 5B, where the values of slope are plotted against the logarithm of number of samples in each dataset. The plot for the second group can tentatively be interpreted to show that larger datasets tend to be characterized by more positive slopes.

In Fig. 5C, data from all datasets in each environment type are combined and used to assess global correlations between Eh7 and *Z*_C_. For Bacteria, the global analysis for each environment type generates a positive slope, but only the Lake & Pond, Groundwater, and Soil groups are deemed significant according to an arbitrary *P*-value cutoff of *<* 0.01. In contrast to Bacteria, Archaea show a negative or no correlation for different environment types at a global scale. Because of the lower number of datasets with archaeal sequences for each environment type, they may not represent true global patterns, and environment types with less than five available datasets are grayed out in Fig. 5C.

Combining data for all environment types except hot springs yields significant positive and negative correlations between Eh7 and *Z*_C_ for Bacteria and Archaea, respectively (Fig. 5D). However, the data for Archaea are dominated by a few environment types (Lake & Pond, Sediment, and Soil); as will be seen below, adding data for hot springs increases the slope of the global linear fit for both Bacteria and Archaea.

### Redox potential and other environmental factors in hot spring communities

Hot spring and river samples in the Copahue Volcano-Río Agrio area in Argentina (Lopez Bedogni et al., 2020; Massello et al., 2020) yield a small negative slope of the linear regression between Eh7 and bacterial *Z*_C_ (Fig. 6A). The other hot spring datasets have some of the highest slopes of linear fits between Eh7 and *Z*_C_ for any environment, but they are also characterized by a wide range of fitted *Z*_C_ between datasets. The environmental factors that contribute to this variation are explored below.

**Fig. 6.**
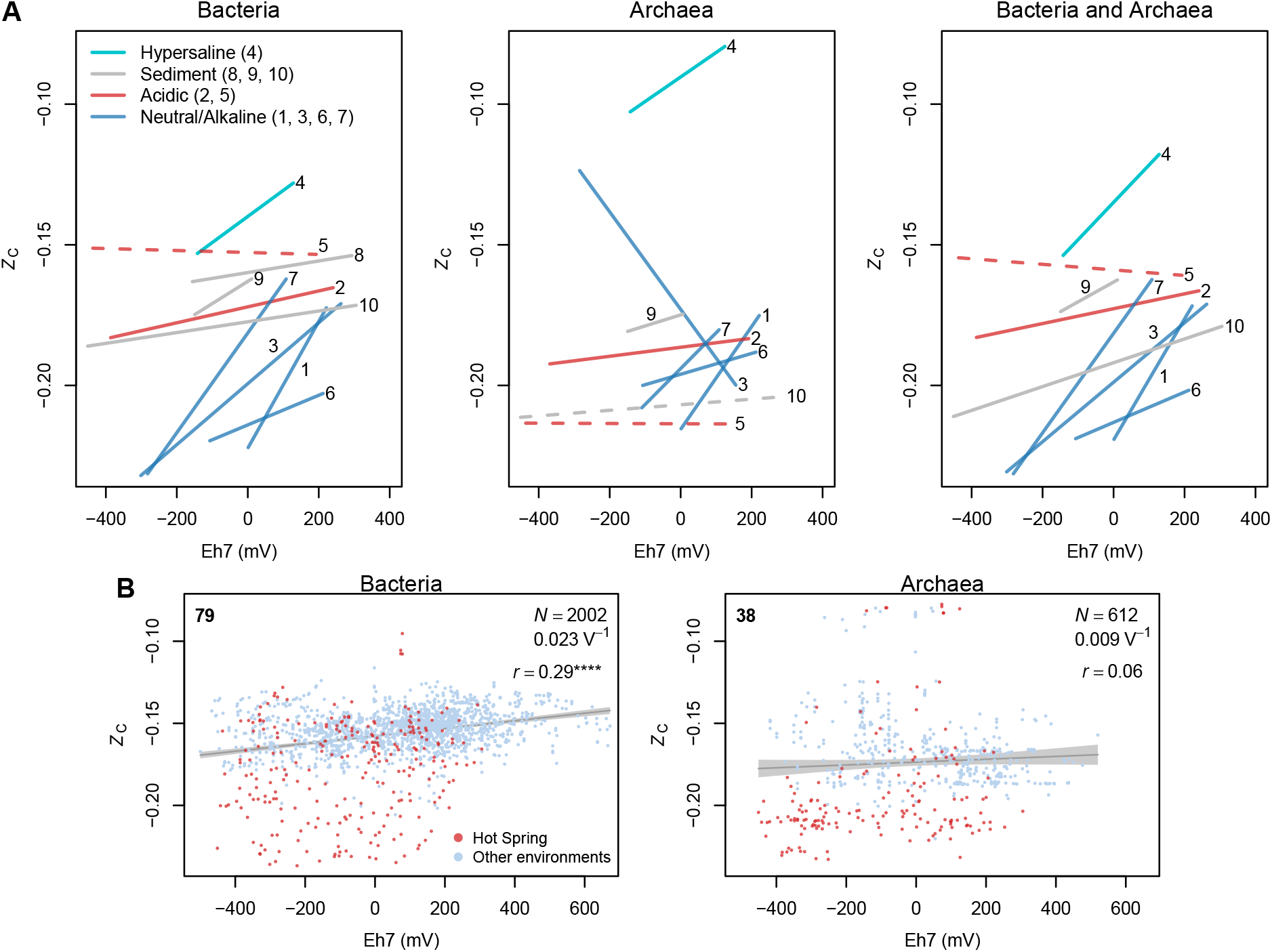
Distinctions in carbon oxidation state estimated for different hot springs, and global fits for all environments. (A) *Z*_C_-Eh7 fits for hot spring datasets using only bacterial abundances, only archaeal abundances, or combined taxonomic abundances for both domains. Colors are used to distinguish major sample types classified as hypersaline water, sediment, acidic water, and neutral to alkaline water. Dashed lines represent fits with a slope of less than 0.01 V^-1^. Numbers for datasets are (1) Bison Pool, Yellowstone National Park (unlike the other datasets, DNA was extracted from biofilm samples) (Swingley et al., 2012), (2) acidic and (3) circumneutral to alkaline New Zealand hot springs (Power et al., 2018), (4) Dallol Geothermal Area (hypersaline and moderate to extremely acidic water) (Belilla et al., 2019), (5) Copahue Volcano-Río Agrio (Lopez Bedogni et al., 2020; Massello et al., 2020), (6) Southwestern Yunnan (seven samples with pH *>* 7 and two moderately acidic samples) (Guo et al., 2021), (7) Eastern Tibetan Plateau (Guo et al., 2020), (8) Uzon Caldera (nine samples for acidic water and sediment and one high-pH sample) (Peltek et al., 2020), (9) Southern Tibetan Plateau (Ma et al., 2021), (10) Japan Hydrothermal Sediment (Omae et al., 2019). See Figure S1 for individual sample values. (B) Global fits for Bacteria and Archaea in all samples from all environment types including hot springs. The legends show (top left) the number of datasets, (top right) the number of samples (*N*), slope of linear regression (units of V^-1^), and Pearson correlation coefficient (*r*) with *P*-value indicated by stars (\*\*\*\**P <* 10^−30^).

A dataset for water samples in the Dallol geothermal area, Kenya (Belilla et al., 2019), has the highest range of fitted *Z*_C_. This area is described as a polyextreme environment that is characterized by hypersaline conditions in addition to high heat and low pH. It is likely that high *Z*_C_ arises not because of higher redox potential, but because of the greater number of acidic residues in proteomes of halophilic Archaea that represent adaptations to stabilize the three-dimensional structure of proteins (Dick et al., 2020). After the dataset for the Dallol geothermal area, those for other acidic springs and sediment samples have somewhat lower ranges of fitted *Z*_C_.

The datasets with the lowest ranges of fitted *Z*_C_ are those for water samples from neutral or alkaline hot springs in Southwestern Yunnan (Guo et al., 2021), the Eastern Tibetan Plateau (Guo et al., 2020) (see also Fig. 4), the circumneutral-to-alkaline subset of New Zealand hot springs (Power et al., 2018), and Bison Pool in Yellowstone National Park, USA (Swingley et al., 2012). The biofilm sampling for DNA analysis of Bison Pool took place in 2005 (Swingley et al., 2012) while redox potential measurements were made in summer of 2009 (Table S1 and Dick and Shock, 2011); Eh measurements are not available for the highest-temperature site of Bison Pool, so a value measured in the neighboring and chemically similar (Boyer et al., 2020) Mound Spring was used. Notably, the range of *Z*_C_ for communities at Bison Pool estimated from 16S rRNA gene sequences in this study (ca. −0.22 to −0.16; Figure S1) largely overlaps with the range inferred from shotgun metagenomic sequencing (ca. −0.21 to −0.155; Dick and Shock, 2011 and Dick et al., 2019), lending support to the accuracy of using estimated community proteomes from 16S rRNA sequencing.

Archaeal abundances in most hot spring datasets yield estimated *Z*_C_ that show a small positive association with Eh7, except for the highly negative slope for circumneutral to alkaline New Zealand hot springs (Fig. 6A). When taxonomic abundances of both domains are combined, an overall positive association between Eh7 and *Z*_C_ remains. This is especially the case for the Dallol water and Japan sediment datasets, where the combined abundances show a higher positive slope than either Archaea or Bacteria alone. This is possible evidence that the relative abundances of the domains themselves, and not only of taxa within the bacterial domain, are chemically shaped by redox conditions in these environments. Additional caution is needed in interpreting the results for the Dallol area because of the likelihood that bacterial sequences in the most acidic samples are contaminants and that archaeal sequences are carried on wind-blown dust (Belilla et al., 2019).

Finally, adding the hot spring data to the global analysis for all environments increases the fitted slope for both Bacteria and Archaea relative to the global correlations without hot springs (compare Figs. 5D and 6B). For both domains, alkaline hot springs have the lowest range of *Z*_C_ compared to other environments. Although non-hydrothermal alkaline springs hosted in ophiolites are also known for very reducing conditions, their range of *Z*_C_ is higher than that for alkaline hot springs (Fig. 5A). It therefore appears that the combination of high temperature, high pH, and reducing conditions in alkaline hot springs may be a crucial combination that selects for organisms with more reduced proteins than any other environment.

## Discussion

Although no taxonomic data were explicitly shown, all of the results described above represent a combination of taxonomic abundances with reference proteomes to estimate a chemical formula for the community proteome of each sample. Because *Z*_C_ at the community level is determined by protein-coding genomic sequences and relative abundances of different organisms, its predicted correlation with redox potential, while being based on thermodynamic concepts, also has theoretical implications for evolution and ecology. More broadly, *Z*_C_ is a quantitative chemical representation of microbial communities that allows predictions about eco-evolutionary differences that are not possible with conventional methods of diversity analysis and sequence comparison.

Our initial results for Winogradsky columns (Fig. 2) demonstrated that *Z*_C_ in microcosm communities decreases, as predicted, as redox potential decreases with depth. Analysis of multiple field-based lake and seawater datasets showed that lower redox potential with depth in stratified water bodies is almost always associated with decreasing *Z*_C_ of the estimated bacterial community proteomes. This mirrors the earlier findings of Dick and Tan (2021) for oxygen gradients in stratified systems. We then performed global comparisons for different environment types considered separately and together (Fig. 5C–D). The positive correlations between Eh7 and *Z*_C_ for Bacteria in various environment types, especially lakes and ponds, groundwater, and soils, and in the pan-environmental comparison provide ample support for the chemical shaping hypothesis.

In contrast to Bacteria, Archaea do not show a strong signal of chemical shaping, and even display a negative correlation for some environment types (Fig. 5C). However, greater uncertainty of archaeal abundances due to primer biases and the occurrence in some environments of some phylogenetic groups, such as Woesearchaeota and Pacearchaeota in the DPANN superphylum (Ortiz-Alvarez and Casamayor, 2016), which are identified by the RDP Classifier but are not currently available in RefSeq, mean that the current findings for Archaea remain provisional. Importantly, hot springs exhibit a wider range of *Z*_C_ than other environment types, and when they are included in the global analysis, the fitted slopes for both Bacteria and Archaea increase, and the correlation for Archaea becomes marginally positive (Fig. 6D). Therefore, compared to other environments, microbial communities in alkaline hot springs may represent the greatest evolutionary and ecological divergence toward more reduced proteins under the influence of reducing conditions.

Multiple physicochemical factors, and not only redox potential, are drivers of microbial community composition in many environments. For instance, salinity gradients have a major influence on microbial community structure in sediments of the Great Salt Lake (GSL), USA (Boyd et al., 2017). The analysis in this study does not control for these other factors. Nevertheless, a positive correlation between Eh7 and *Z*_C_ for both Archaea and Bacteria in the GSL sediment dataset (Figure S1) speaks to the potential for detecting a signal of redox potential from among other environmental drivers. The positive slopes of linear regressions for many datasets from aquatic and soil environments (Fig. 5B) are consistent with the prediction of local chemical shaping by redox potential, which may be superimposed on the effects of other environmental factors. This is particularly evident for different hot spring datasets, which are offset from one another due to differences in salinity, pH, and sample type, but with one exception exhibit positive correlations between Eh7 and bacterial *Z*_C_ at the local scale (Fig. 6).

A chemical representation of microbial communities naturally suggests a strategy to model deterministic outcomes that can be predicted by thermodynamics. This type of assessment might complement approaches that depend on variation partitioning or null models to discern stochastic and deterministic community assembly processes (Stegen et al., 2015) but do not explicitly account for chemical variability. Although the influence of ecological determinism is thought to scale with body size, so that microorganisms display relatively low degrees of determinism in community assembly (Farjalla et al., 2012; Luan et al., 2020), the results of this study suggest that numerous bacterial communities are chemically shaped to some extent through a previously unrecognized nexus between environment and proteome composition.

## Declarations

### Data and code availability

All data analyzed in this study are from public sources; see Table 1 for accession numbers for sequence data and Table S1 for sources of redox potential values. All data and code generated in this study are in the JMDplots R package that is available on GitHub (https://github.com/jedick/JMDplots); see Section Computer code for data analysis and visualization for details.

### Supplementary material

Table S1 Dataset summaries: Bibliographic study key, name, number of samples with bacterial and archaeal sequences, sample type (soil, sediment, water, or biological), ranges of *T*, pH, Eh, and Eh7, sign of *Z*_C_–Eh7 correlation for Bacteria, source of Eh data, notes.

Table S2 Sequence processing statistics.

Figure S1 *Z*_C_–Eh7 scatterplots and linear regressions for all datasets.

Dataset S1 Data and metadata for each sample: Study key, environment type, lineage (Bacteria or Archaea), sample name, SRA or MG-RAST accession number, grouping description, sample group, values of Eh, Eh7, and *Z*_C_.

Dataset S2 Coefficients of linear regressions and Eh7 ranges for each dataset and domain. Figure S1 and Datasets S1 and S2 were generated using the `orp16S S1()` function in the JMDplots package.

Dataset S3 BibTeX bib file for data sources.

