## Supplementary material for "Local and global chemical shaping of bacterial communities by redox potential": Table S2

**Table S2.** Sequence processing statistsics.

| Study | BioProject | Samples | Available reads per sample(a) | Filtered reads per sample(b) | Filtered read length | Sing. %(c) | Chim. %(d) | Reads per sample used for classification | % Classification to genus level | % Mapping to NCBI taxonomy |
|---|---|---|---|---|---|---|---|---|---|---|
| **River & Seawater** | | | | | | | | | | |
| MLL+18 | PRJNA319446 PRJNA357334 | 47 | 37842 | 35912 | 250 (i) | 19.9 | 0.8 | 9923 | 39 | 96.7 |
| SVH+19 | PRJNA423140 | 15 | 6631 | 5604 | (f) | 20.5 | 7.2 | 4132 | 17 | 98.3 |
| HXZ+20 | PRJNA503500 | 21 (v) | 92727 | 75235 | 440 | 34.3 | 2.3 | 9773 | 44 | 95.8 |
| KLY+20 | PRJNA530625 PRJNA594996 | 17 | 80103 | 54187 | 250 (i) | 27.3 | 12.8 | 8723 | 63 | 78.2 |
| WHL+21 | PRJNA588356 | 52 | 79313 | 57128 | 400 | 61.8 | 3.9 | 9555 | 62 | 96.2 |
| LXH+20 | PRJNA636956 | 17 (n) | 55689 | 33760 | 450 (i) | 65.3 | 4.7 | 8025 | 61 | 97.5 |
| JVW+20 | PRJNA656714 | 17 | 189897 | 29005 | 250 | 34.9 | 6.9 | 8948 | 38 | 95.2 |
| ZZL+21 | PRJNA681688 | 54 | 51033 | 35179 | 450 | 41.7 | 8.5 | 8753 | 43 | 93.3 |
| GZL21 | PRJNA733826 | 94 | 45674 | 19625 | 420 | 83.0 | 8.7 | 2966 | 53 | 93.7 |
| **Lake & Pond** | | | | | | | | | | |
| SAR+13 | PRJNA174394 | 50 (k) | 3939 | 2686 | (g) | (h) | 5.1 | 2549 | 23 | 87.7 |
| LZR+17 | PRJNA255556 | 11 | 161057 | 157861 | 250 (i) | 12.4 | 4.5 | 9549 | 68 | 96.4 |
| ECS+18 | PRJNA289691 | 8 | 95842 | 60821 | 400 (i) (t) | 39.2 | 8.2 | 9180 | 32 | 99.9 |
| LLC+19 | PRJNA315049 | 36 | 74052 | 52599 | 400 (i) | 19.0 | 7.6 | 9245 | 32 | 94.3 |
| SCH+16 | PRJNA321351 | 9 | 1029567 (w) | 89661 | 250 (j) | 17.4 | 9.9 | 9010 | 31 | 91.8 |
| BCA+21 | PRJNA395513 | 36 | 280205 | 95996 | 450 | 34.9 | 9.9 | 8638 | 44 | 97.9 |
| HLZ+18 | PRJNA407260 | 27 | 88141 | 82514 | 420 | 32.2 | 5.8 | 9415 | 48 | 95.2 |
| CNA+20 | PRJNA473136 | 15 | 81278 | 80204 | 450 | 26.6 | 10.0 | 9004 | 49 | 92.1 |
| BWD+19 | PRJNA489447 | 102 (o) | 76580 | 63910 | 400 | 25.8 | 23.5 | 6900 | 43 | 98.6 |
| RSJ+21 | PRJNA516525 | 38 | 184093 | 112153 | 290 | 14.5 | 3.7 | 9596 | 39 | 91.3 |
| BOEM21 | PRJNA641339 | 9 | 41575 | 35832 | 400 | 37.3 | 5.0 | 9504 | 55 | 87.4 |
| GSY+20 | PRJEB38636 | 19 | 138806 | 121640 | 120 (j) | 1.8 | 0.2 | 9981 | 51 | 93.7 |
| NLE+21 | PRJEB39923 | 21 | 145541 | 110098 | 440 | 48.9 | 2.5 | 9746 | 51 | 97.3 |
| FAV+21 | PRJNA731062 | 17 | 245634 | 74601 | 400 | 30.3 | 12.4 | 9412 | 44 | 90.7 |
| GRG+20 | mgp83146 | 6 | 63388 | 63388 | (y) | 33.8 | 7.9 | 9211 | 43 | 88.6 |
| **Hot Spring** | | | | | | | | | | |
| SMS+12 | NA (from authors) | 5 | 464 | 464 | (u) | (u) | (u) | 464 | 77 | 75.8 |
| PCL+18 | PRJEB24353 | 81 (p) | 43631 | 24507 | 250 (q) | 24.9 | 2.6 | 8863 | 62 | 91.5 |
| BMJ+19 | PRJNA541281 | 22 | 173978 | 138702 | 300 | 37.0 | 7.0 | 9302 | 62 | 94.3 |
| LMG+20 | PRJNA542136 | 9 | 188747 | 109074 | 280 (j) | 30.8 | 9.4 | 9057 | 73 | 96.8 |
| MCC+20 | | | | | | | | | | |
| GWSS21 | PRJNA548851 | 9 | 36256 | 29951 | 300 | 15.1 | 3.7 | 9627 | 79 | 98.4 |
| GWS+20 | PRJNA592622 | 16 | 30006 | 28946 | 300 | 17.2 | 5.8 | 9416 | 44 | 98.3 |
| PBU+20 | PRJNA623081 | 10 | 34728 | 15016 | 400 | 67.7 | 11.6 | 4849 | 86 | 96.8 |
| MWY+21 | PRJNA638734 | 13 | 58142 | 53076 | 250 | 37.2 | 5.0 | 8996 | 39 | 90.3 |
| OFY+19 | NA (OTU Table) | NA | NA | NA | NA | NA | NA | NA | 23 | 75.9 |
| **Alkaline Spring** | | | | | | | | | | |
| SBP+20 | PRJNA289273 | 24 | 82213 | 67456 | 250 | 8.4 | 14.3 | 8571 | 57 | 95.5 |
| RMB+17 | PRJNA352492 | 20 | 49966 | 44726 | 250 | 19.2 | 2.4 | 9764 | 37 | 95.0 |
| CTS+17 | PRJNA361138 | 66 | 82772 | 71957 | 400 | 77.1 | 10.5 | 6331 | 35 | 98.8 |
| KSR+21 | PRJNA685937 | 8 | 73796 | 60037 | 440 | 35.5 | 11.6 | 8844 | 25 | 92.3 |
| NTB+21 | PRJNA743134 | 37 | 23417 | 19081 | 150(j) | 12.1 | 8.2 | 5041 | 32 | 99.8 |
| **Groundwater** | | | | | | | | | | |
| KLM+16 | PRJNA308958 | 30 | 9917 | 6076 | (f) | 58.4 | 3.0 | 2455 | 43 | 91.5 |
| YHK+19 | PRJDB5959 | 13 | 91096 | 25089 | 250 (i) (x) | 37.6 | 1.1 | 9801 | 37 | 97.2 |
| SDH+19 | PRJNA408058 | 6 | 37333 | 36503 | 300 | 53.8 | 11.6 | 8839 | 65 | 99.3 |
| SRM+19 | PRJNA434769 | 19 | 154589 | 30740 | 250 | 31.2 | 11.7 | 8426 | 47 | 93.5 |
| APV+20 | PRJNA488796 | 24 | 86279 | 69012 | 250 | 25.3 | 7.2 | 9094 | 55 | 90.7 |
| SKP+21 | PRJNA528471 | 12 | 98161 | 11218 | 200 | 38.1 | 5.5 | 5910 | 19 | 96.5 |
| YHK+20 | PRJEB33032 | 33 (l) | 99926 | 48564 | 400 | 39.6 | 6.8 | 7412 | 33 | 55.2 |
| JDP+20 | PRJNA553521 | 8 | 188240 | 135572 | 275 (i) | 19.6 | 14.8 | 8524 | 65 | 99.9 |
| GWS+19 | PRJNA623081 | 11 | 31412 | 33576 | 300 | 13.7 | 5.4 | 9464 | 35 | 99.1 |
| SRM+21 | PRJNA630252 | 13 | 87292 | 36180 | 250 | 37.6 | 12.8 | 8718 | 34 | 94.2 |
| ZCZ+21 | PRJNA667833 | 18 | 22010 | 19704 | 450 | (m) | 49.9 | 4721 | 54 | 95.0 |
| **Sediment** | | | | | | | | | | |
| JHL+12 | SRP009131 | 15 | 5639 | 3566 | (f) | 35.6 | 5.0 | 2183 | 54 | 99.2 |
| GFE+16 | PRJNA262691 | 16 | 25170 | 21905 | (s) | 65.8 | 33.7 | 4557 | 16 | 89.8 |
| ZML+17 | PRJEB12432 PRJEB12429 | 66 | 25188 | 19726 | 400 (i) | 77.6 | 8.9 | 3945 | 38 | 97.8 |
| BYB+17 | PRJNA319444 | 16 | 13652 | 7189 | (f) | (h) | 22.4 | 5577 | 38 | 93.1 |
| BSPD17 | PRJNA322450 | 32 | 97803 | 49495 | 400 | 78.7 | 6.7 | 5386 | 75 | 86.1 |
| HDZ+19 | PRJNA352457 | 28 | 158622 | 97143 | 420 | 52.2 | 6.2 | 9380 | 55 | 92.2 |
| WHLH21 | PRJNA400089 | 20 | 60168 | 16758 | 400 (i) | 38.9 | 0.8 | 7927 | 47 | 95.6 |
| SCM+18 | PRJNA429278 | 11 | 144324 | 52805 | 250 (j) | 47.8 | 10.4 | 8962 | 22 | 97.2 |
| CLS+19 | PRJNA437688 PRJNA437695 PRJNA437692 PRJNA437697 | 45 | 145210 | 138438 | 150 (i) | 69.4 | 0.1 | 8812 | 33 | 94.5 |
| ZDA+20 | PRJEB28365 | 16 | 69934 | 51796 | 450 (i) | 62.0 | 10.5 | 8798 | 58 | 89.0 |
| VMB+19 | PRJNA453733 | 6 | 106023 | 68644 | 380 (i) | 47.8 | 3.1 | 9687 | 54 | 92.2 |
| WHC+19 | PRJNA488529 | 12 | 122751 | 101233 | 420 | 54.9 | 8.6 | 9236 | 80 | 91.2 |
| HSF+19 | PRJNA507590 | 63 | 47969 | 31059 | 440 | 35.2 | 4.3 | 8370 | 30 | 84.3 |
| MCS+21 | PRJEB31703 | 14 | 101917 | 75549 | 400 | 61.7 | 2.6 | 7893 | 35 | 95.1 |
| LMBA21 | PRJEB35647 | 256 | 63613 | 43031 | 280 (j) | 52.3 | 8.4 | 8149 | 36 | 93.6 |
| ZZLL21 | PRJNA616197 | 71 | 58505 | 38640 | 450 | 60.3 | 10.4 | 7303 | 53 | 88.3 |
| WFB+21 | PRJNA639965 | 83 | 115656 | 87532 | 440 | 32.9 | 4.7 | 8936 | 35 | 88.6 |
| **Soil** | | | | | | | | | | |
| SBW+17 | PRJNA341915 | 16 | 52013 | 36856 | 280 (j) | 52.4 | 15.7 | 8428 | 34 | 70.3 |
| MLL+19 | PRJNA361046 | 30 | 43372 | 36941 | 250 (i) | 46.1 | 1.7 | 9509 | 32 | 83.8 |
| BMOB18 | PRJNA415514 | 48 | 13229 | 11791 | 300 (i) | 41.0 | 9.7 | 5937 | 23 | 89.1 |
| ZZZ+18 | PRJNA427749 PRJNA427853 | 36 | 43487 | 38493 | 240 (j) | 16.3 | 4.6 | 9430 | 67 | 95.7 |
| PMM+20 | PRJEB28313 | 6 | 53965 | 34921 | 400 | 80.5 | 29.0 | 4825 | 48 | 80.2 |
| ZHZ+19 | PRJNA523725 | 31 | 132508 | 126589 | 300 | 77.9 | 7.7 | 9230 | 23 | 87.7 |
| CWC+20 | PRJNA564714 | 84 | 83475 | 68477 | 440 | 52.2 | 8.2 | 9183 | 62 | 72.8 |
| PSG+20 | PRJNA575041 | 49 | 77866 | 21021 | 250 | 29.7 | 2.9 | 9435 | 55 | 57.6 |
| XLD+20 | PRJNA576993 | 48 | 27908 | 24913 | 250 (i) | 38.2 | 4.5 | 8351 | 71 | 87.6 |
| LJC+20 | PRJNA607877 | 45 | 57625 | 53602 | 420 | 79.0 | 3.9 | 8542 | 50 | 91.8 |
| DTJ+20 | PRJNA611687 | 36 | 51462 | 34005 | 420 | 79.8 | 7.3 | 6392 | 41 | 80.0 |
| LLL+21 | PRJNA665924 | 15 | 56756 | 36987 | 450 | 70.4 | 9.5 | 7134 | 51 | 79.4 |
| DLS21 | PRJNA690162 | 143 | 131364 | 79634 | 300 (j) | 97.5 | 2.0 | 1909 | 35 | 74.4 |
| **Winogradsky Columns** | | | | | | | | | | |
| RBW+14 | PRJNA234104 | 53 (r) | 12933 | 7507 | 250 | 44.9 | 2.6 | 4032 | 47 | 84.4 |

a. Paired forward and reverse reads are counted as one. Paired-end reads were merged with "vsearch -fastq_mergepairs" with default options; reads that failed merging were excluded from the subsequent analysis. All "reads per sample" columns are averages of all samples.

b. Quality filtering was done with "vsearch -fastq_filter" with the options "-fastq_trunclen *length* -fastq-qmax 41 -fastq_maxee_rate 0.005" with length value given in next column.

c. Reads in all samples (runs) were pooled and singletons (sequences appearing exactly once) were identified with "vsearch -derep_fulllength" with the option "-maxuniquesize 1". The singletons were removed from the pooled file using "seqtk subseq"; the remaining sequences for each run were extracted from the pooled file using an awk command. After removing the singletons, for runs with more than 10000 reads, 10000 reads were subsampled using "vsearch -fastx_subsample" with the options "-sample_size 10000" and "-randseed 1234".

d. After subsampling, runs were pooled again and chimeras were removed using "vsearch -uchime_ref" with the option "-nonchimeras" to output non-chimeric sequences (i.e. those not classified as either chimeras or borderline chimeras). The remaining sequences for each run were extracted from the output and used for taxonomic classification.

f. For these 454 sequencing experiments, filtering was done with "vsearch -fastq_filter" with options "-fastq_minlen 200 -fastq_maxlen 600 -fastq_truncqual 15" for read length and quality filtering.

g. As above, but with -fastq_truncqual 11.

h. Because of the high number of singletons and low sequence count, singletons were not removed.

i. Only forward reads available.

j. Only forward reads used.

k. This is number of runs (there are multiple runs for most samples).

l. Excludes 5 samples with failed merging.

m. A large majority of sequences were identified as singletons, so they were not removed prior to subsampling.

n. Only the first available replicate for each sample was used.

o. Only one replicate with the most runs for each sample was used.

p. Randomly selected 100 SRA runs, kept samples with available ORP values on the 1000Springs website, and filtered remaining samples into acidic (42 with $pH < 3$) and neutral-alkaline (39 with $pH > 6$) groups.

q. Only forward reads available; for these Ion Torrent PGM sequences, the option -fastq_qmax 45 was added.

r. Only includes runs listed in SI Table of paper and excludes one run (SRR1140937) with low number of spots (1).

s. As in (f), but with -fastq_truncqual 11 and -fastq_minlen 100.

t. Used -fastq_qmax 93.

u. Because only FASTA files are available, no quality filtering was done; also, because of the very low numbers of reads, no singleton or chimera removal was done.

v. Sequences from Station C4 were used for samples outside the Blue Hole.

w. Because of the large number of available reads, only the first 100000 from each run were processed.

x. Use -fastq_maxee_rate 0.05 (not the 0.005 used for other datasets).

y. Because only FASTA files are available, no quality filtering was done.
