## Supplementary figures and images for "Local and global chemical shaping of bacterial communities by redox potential"

### Figure S1

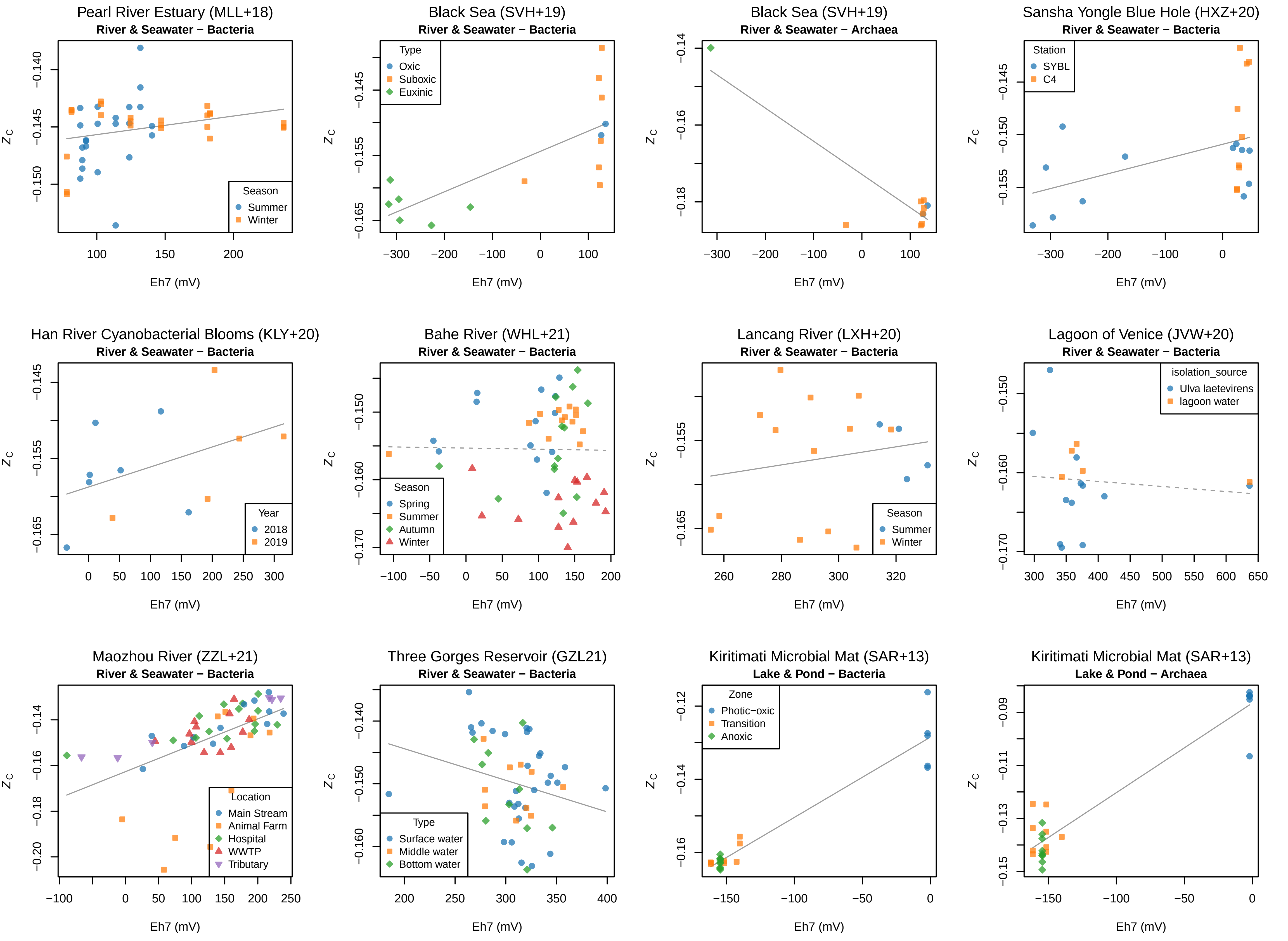

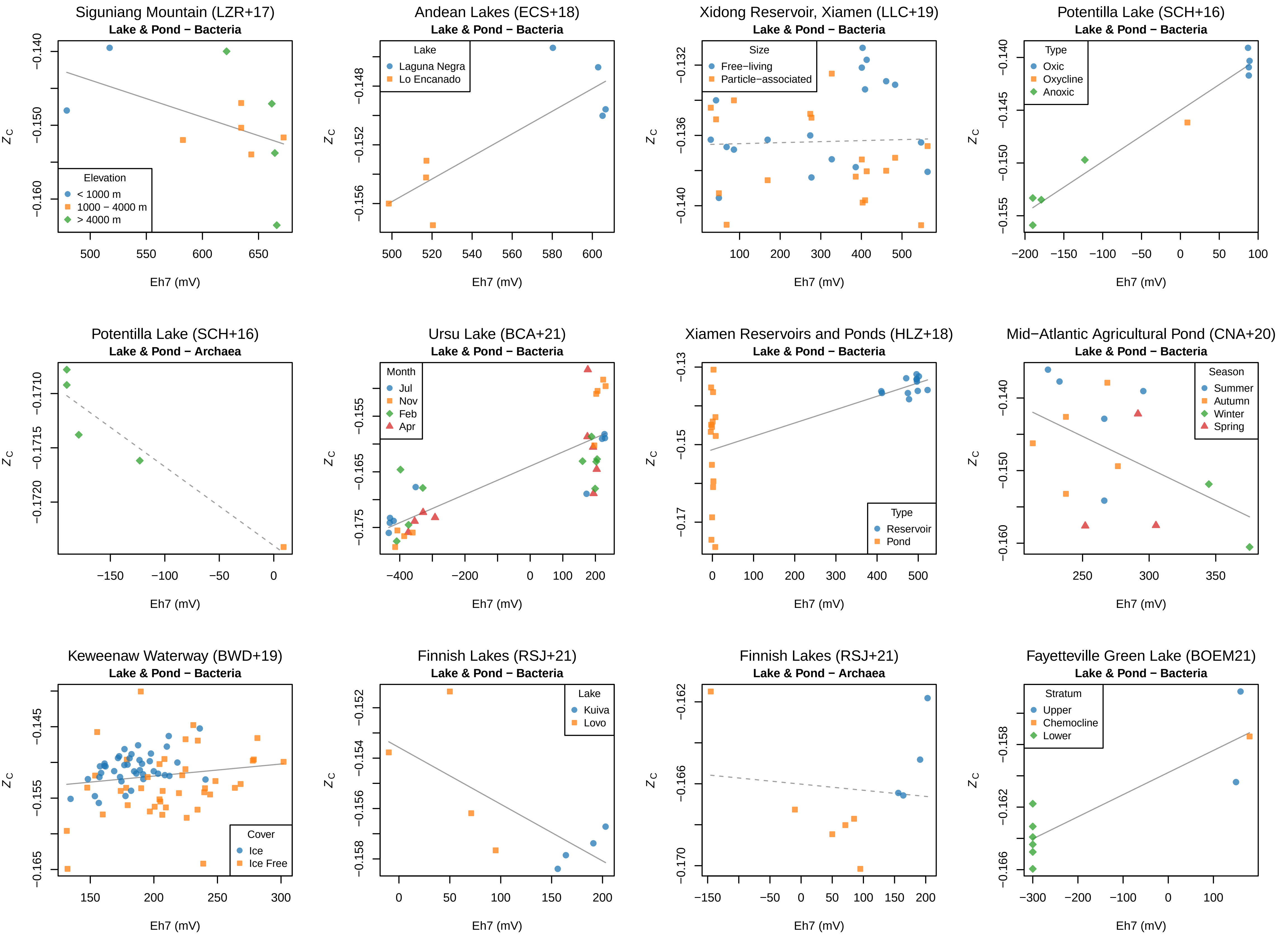

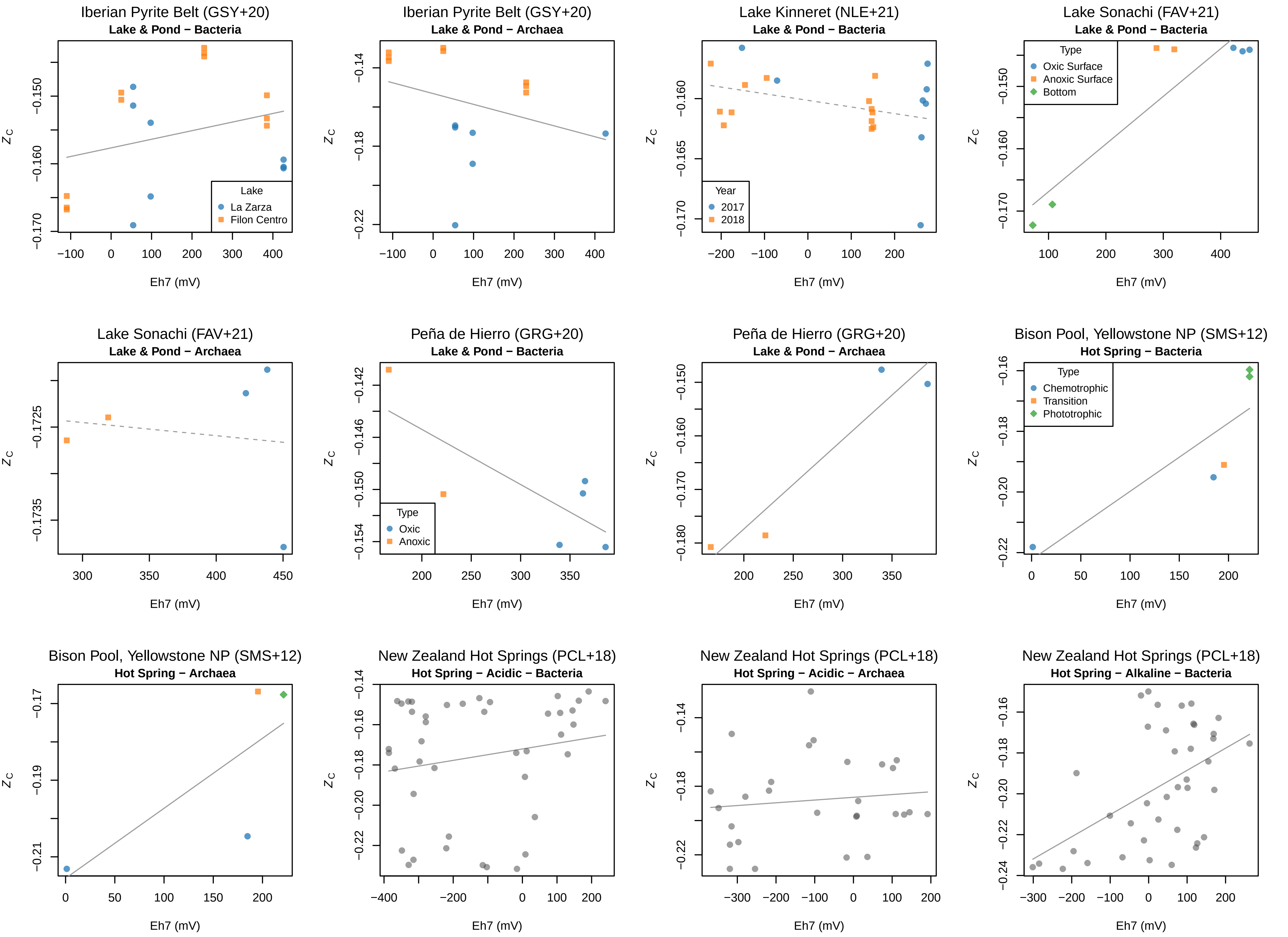

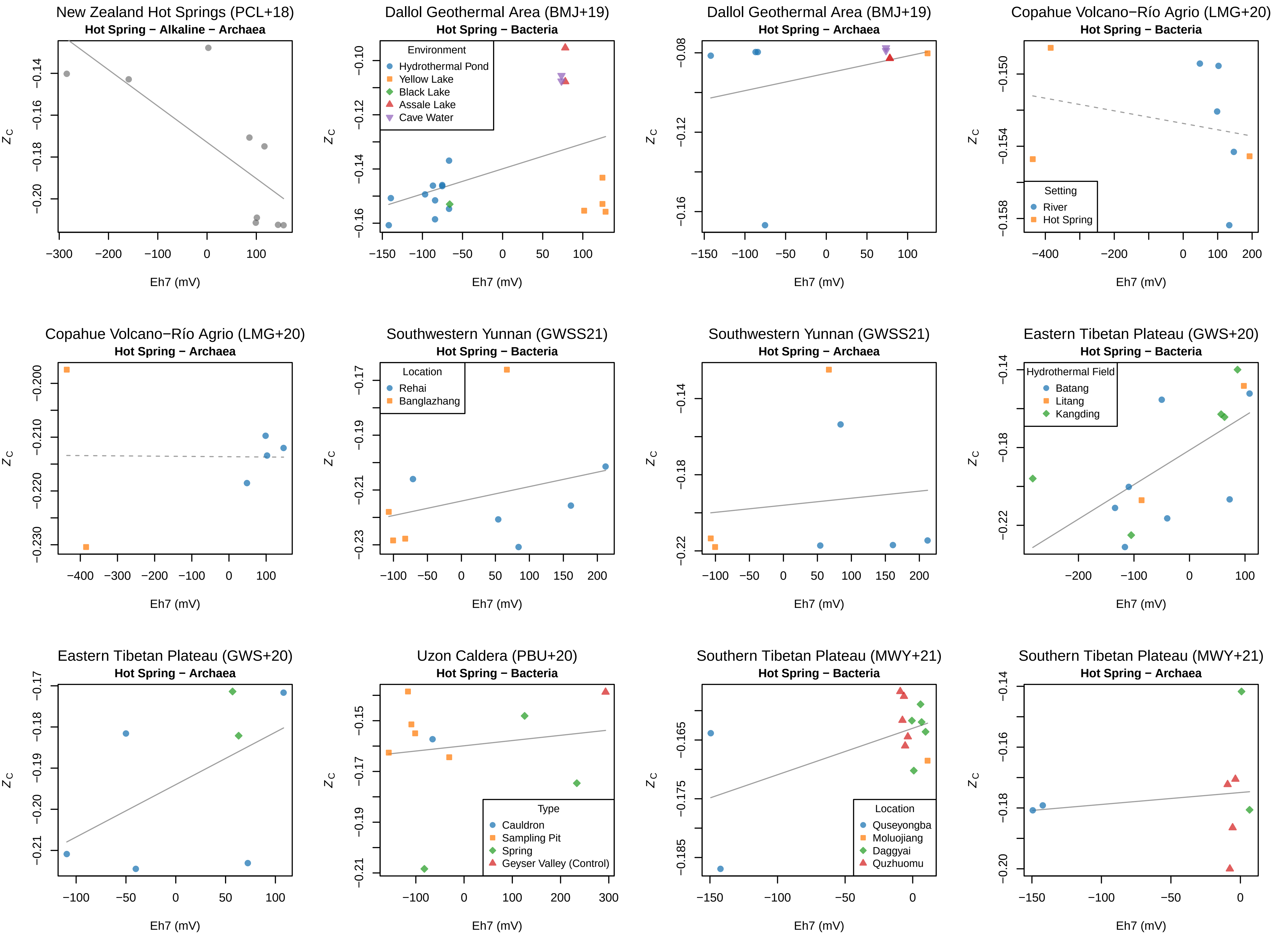

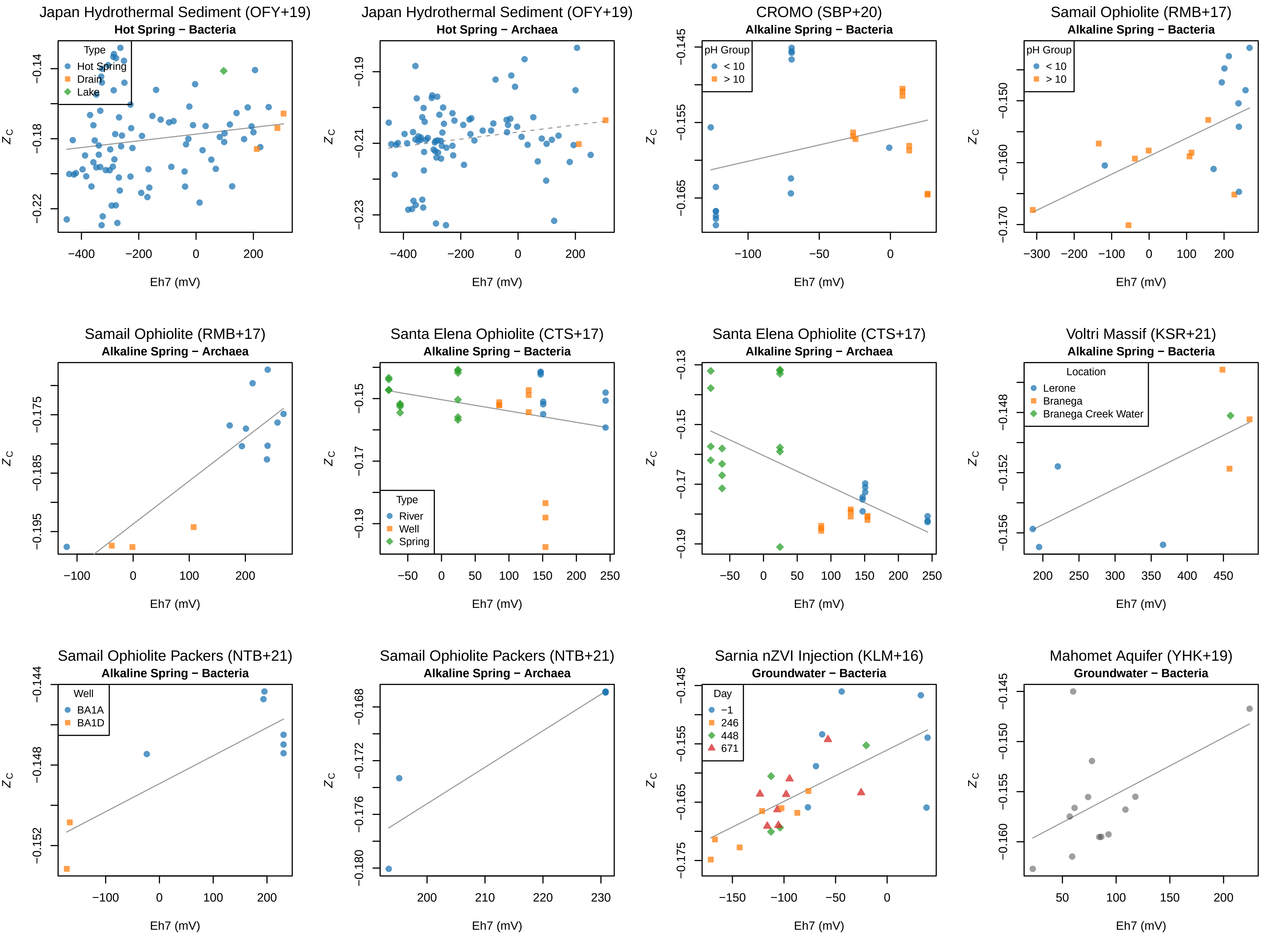

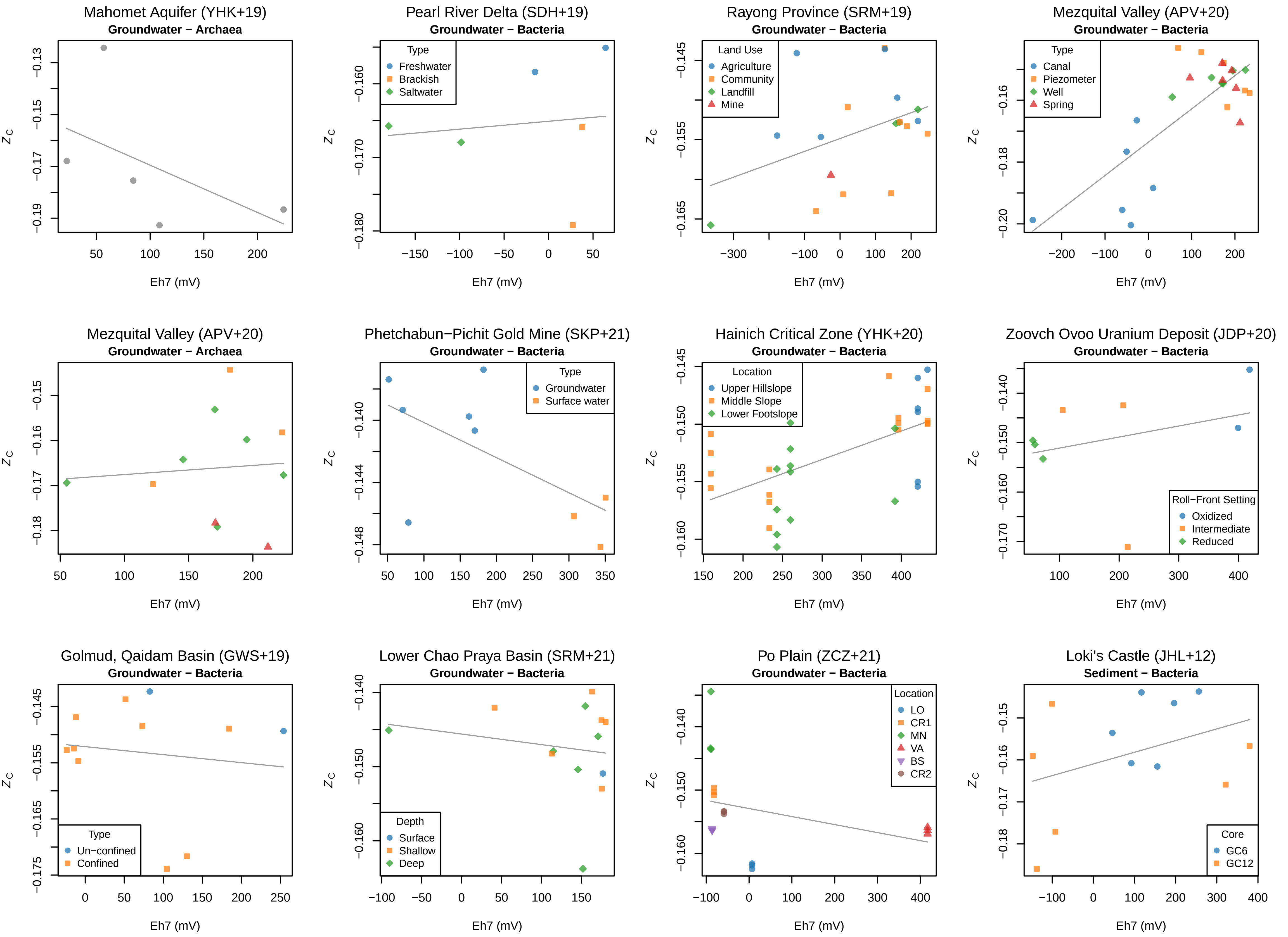

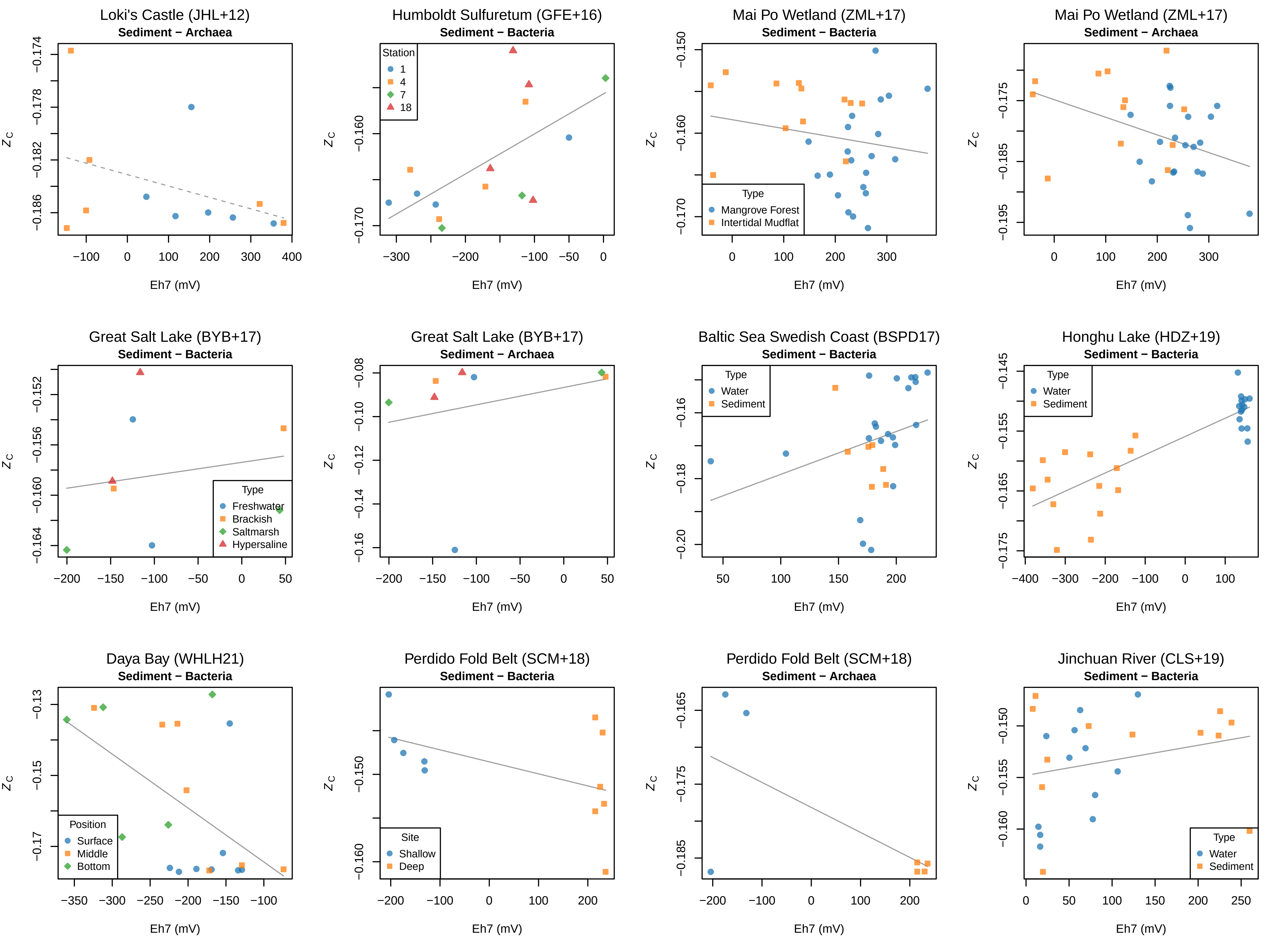

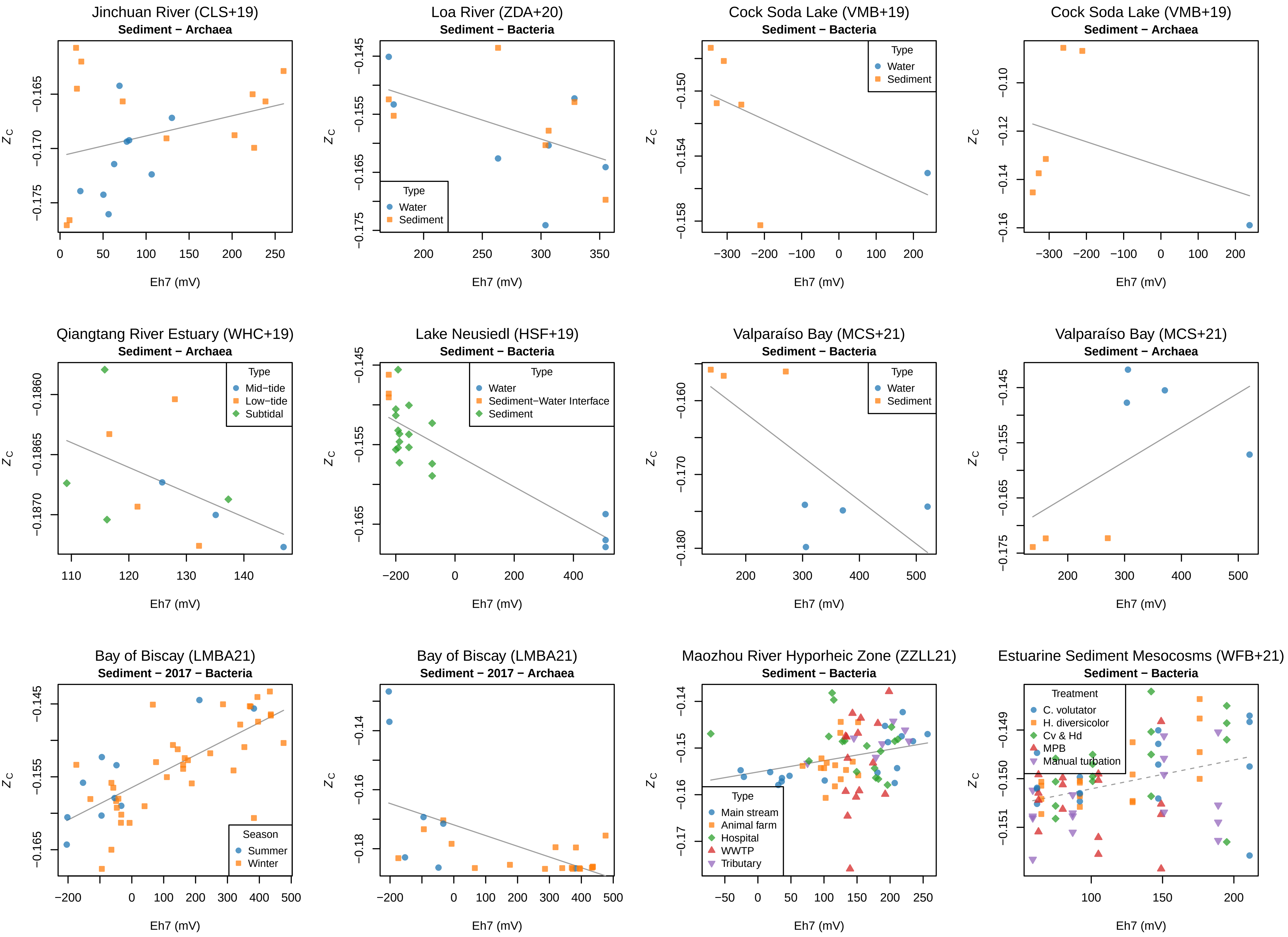

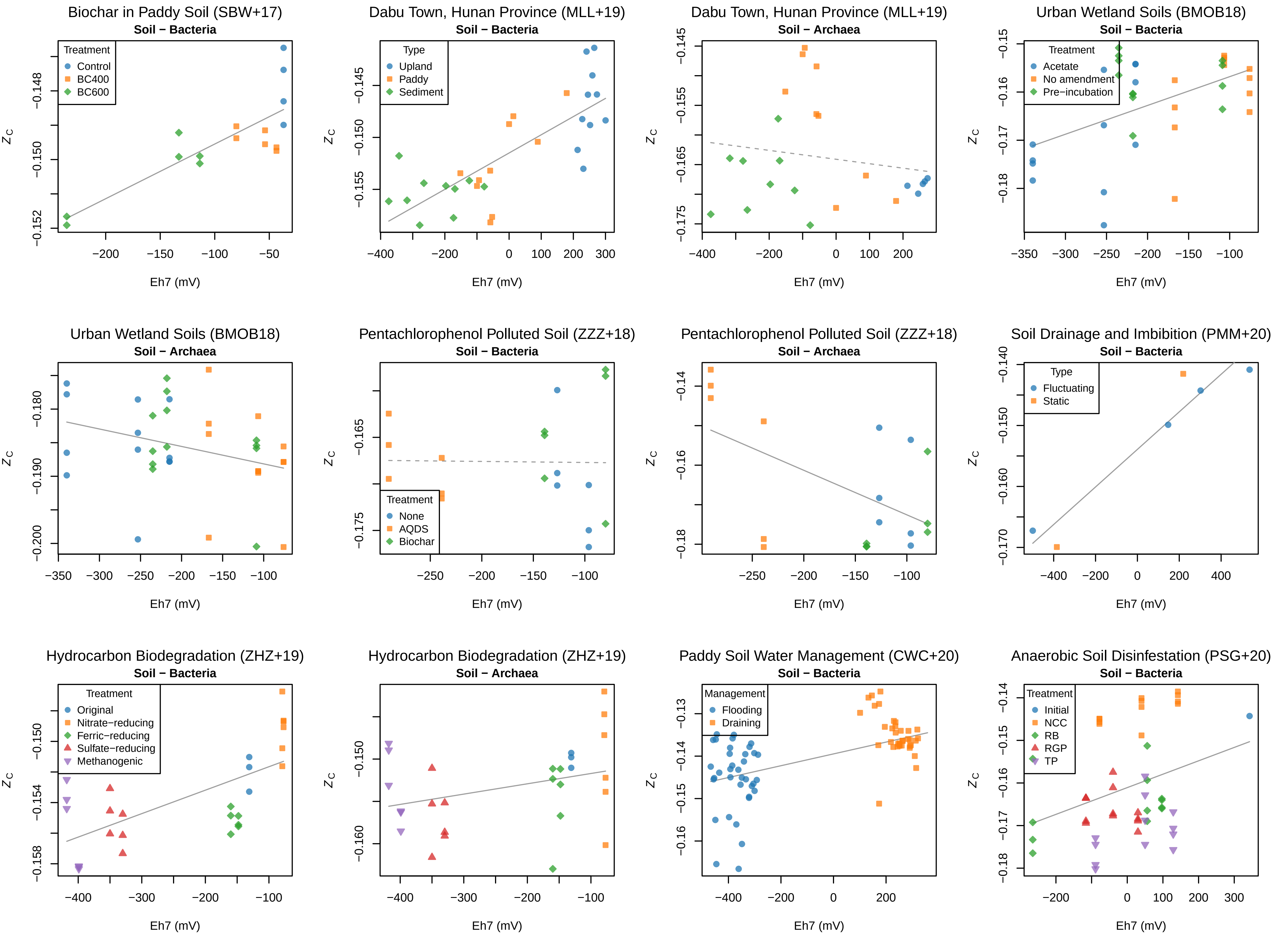

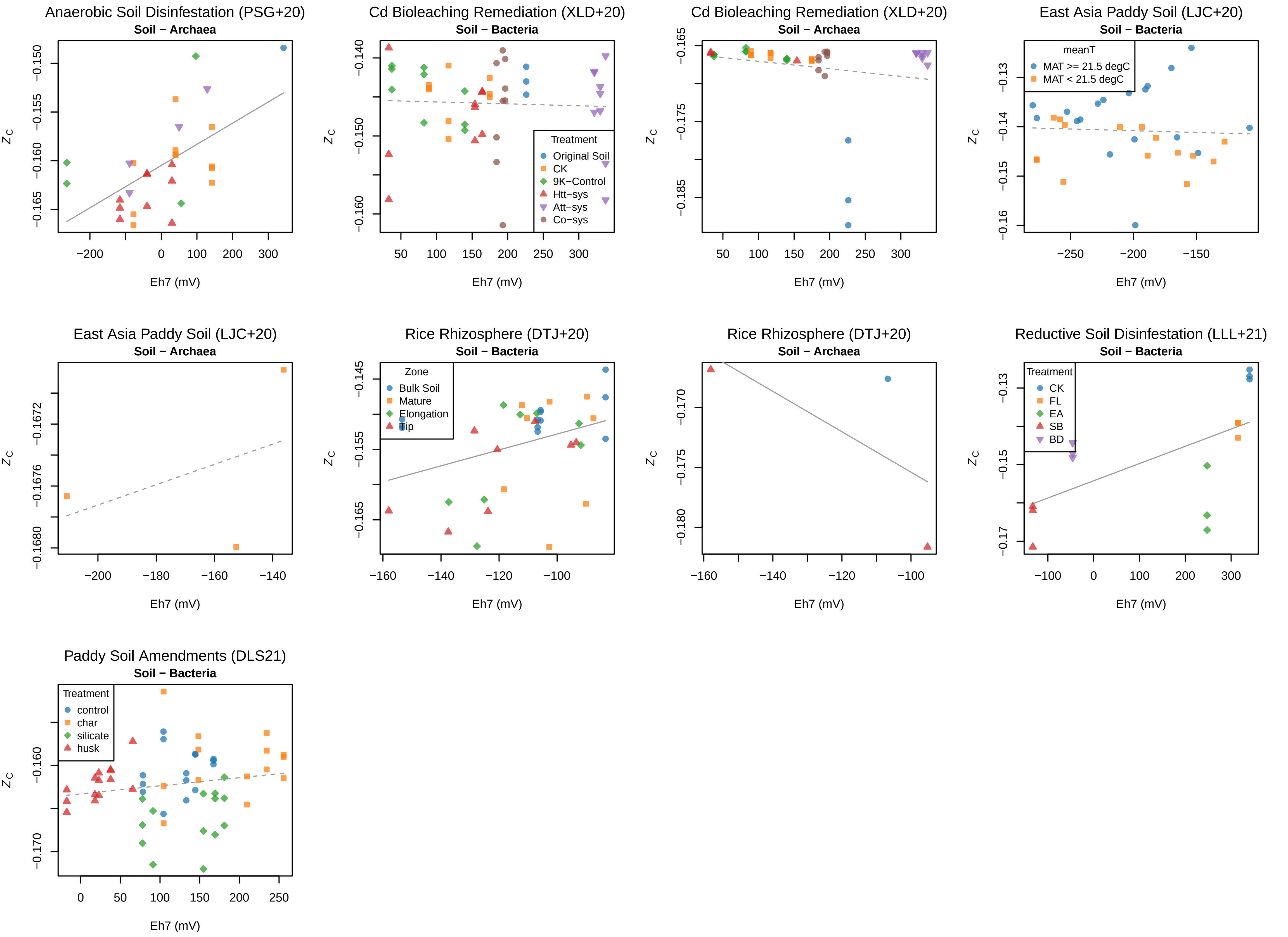
