## Supplementary material for "Local and global chemical shaping of bacterial communities by redox potential": Dataset S3

@Article{DS11,
  author    = {Dick, Jeffrey M. and Shock, Everett L.},
  journal   = {PLOS One},
  title     = {Calculation of the relative chemical stabilities of proteins as a function of temperature and redox chemistry in a hot spring},
  year      = {2011},
  number    = {8},
  pages     = {e22782},
  volume    = {6},
  doi       = {10.1371/journal.pone.0022782},
}

@Article{SMS+12,
  author    = {Swingley, Wesley D. and Meyer-Dombard, D'Arcy R. and Shock, Everett L. and Alsop, Eric B. and Falenski, Heinz D. and Havig, Jeff R. and Raymond, Jason},
  journal   = {PLOS One},
  title     = {{C}oordinating environmental genomics and geochemistry reveals metabolic transitions in a hot spring ecosystem},
  year      = {2012},
  number    = {6},
  pages     = {e38108},
  volume    = {7},
  doi       = {10.1371/journal.pone.0038108},
}

@Article{SAR+13,
  author    = {Schneider, Dominik and Arp, Gernot and Reimer, Andreas and Reitner, Joachim and Daniel, Rolf},
  journal   = {PLOS One},
  title     = {Phylogenetic analysis of a microbialite-forming microbial mat from a hypersaline lake of the {K}iritimati atoll, {C}entral {P}acific},
  year      = {2013},
  number    = {6},
  pages     = {1--14},
  volume    = {8},
  doi       = {10.1371/journal.pone.0066662},
}

@Article{SCH+16,
  author    = {Schütte, Ursel M. E. and Cadieux, Sarah B. and Hemmerich, Chris and Pratt, Lisa M. and White, Jeffrey R.},
  journal   = {Frontiers in Microbiology},
  title     = {Unanticipated geochemical and microbial community structure under seasonal ice cover in a dilute, dimictic {A}rctic lake},
  year      = {2016},
  pages     = {1035},
  volume    = {7},
  doi       = {10.3389/fmicb.2016.01035},
}

@Article{BSPD17,
  author    = {Broman, Elias and Sj{\"o}stedt, Johanna and Pinhassi, Jarone and Dopson, Mark},
  journal   = {Microbiome},
  title     = {Shifts in coastal sediment oxygenation cause pronounced changes in microbial community composition and associated metabolism},
  year      = {2017},
  number    = {1},
  pages     = {96},
  volume    = {5},
  doi       = {10.1186/s40168-017-0311-5},
}

@Article{JHL+12,
  author    = {Jorgensen, Steffen Leth and Hannisdal, Bjarte and Lanzen, Anders and Baumberger, Tamara and Flesland, Kristin and Fonseca, Rita and Ovreas, Lise and Steen, Ida H. and Thorseth, Ingunn H. and Pedersen, Rolf B. and Schleper, Christa},
  journal   = {Proceedings of the National Academy of Sciences},
  title     = {Correlating microbial community profiles with geochemical data in highly stratified sediments from the {A}rctic {M}id-{O}cean {R}idge},
  year      = {2012},
  number    = {42},
  pages     = {E2846--E2855},
  volume    = {109},
  doi       = {10.1073/pnas.1207574109},
}

@Article{ECS+18,
  author    = {Echeverría-Vega, Alex and Chong, Guillermo and Serrano, Antonio E. and Guajardo, Mariela and Encalada, Olga and Parro, Victor and Blanco, Yolanda and Rivas, Luis and Rose, Kevin C. and Moreno-Paz, Mercedes and Luque, José A. and Cabrol, Nathalie A. and Demergasso, Cecilia S.},
  journal   = {Frontiers in Microbiology},
  title     = {{W}atershed-induced limnological and microbial status in two oligotrophic {A}ndean lakes exposed to the same climatic scenario},
  year      = {2018},
  pages     = {357},
  volume    = {9},
  doi       = {10.3389/fmicb.2018.00357},
}

@Article{VMB+19,
  author    = {Vavourakis, Charlotte D. and Mehrshad, Maliheh and Balkema, Cherel and van Hall, Rutger and Andrei, Adrian-{\c S}tefan and Ghai, Rohit and Sorokin, Dimitry Y. and Muyzer, Gerard},
  journal   = {BMC Biology},
  title     = {Metagenomes and metatranscriptomes shed new light on the microbial-mediated sulfur cycle in a {S}iberian soda lake},
  year      = {2019},
  number    = {1},
  pages     = {69},
  volume    = {17},
  doi       = {10.1186/s12915-019-0688-7},
}

@Article{GWS+20,
  author    = {Liang Guo and Guangcai Wang and Yizhi Sheng and Xiaoyi Sun and Zheming Shi and Qingyu Xu and Wenqing Mu},
  journal   = {Science of The Total Environment},
  title     = {Temperature governs the distribution of hot spring microbial community in three hydrothermal fields, {E}astern {T}ibetan {P}lateau {G}eothermal {B}elt, {W}estern {C}hina},
  year      = {2020},
  pages     = {137574},
  volume    = {720},
  doi       = {10.1016/j.scitotenv.2020.137574},
}

@Article{PSG+20,
  author    = {Poret-Peterson, Amisha T. and Sayed, Nada and Glyzewski, Nathaniel and Forbes, Holly and González-Orta, Enid T. and Kluepfel, Daniel A.},
  journal   = {Microbial Ecology},
  title     = {Temporal responses of microbial communities to anaerobic soil disinfestation},
  year      = {2020},
  number    = {1},
  pages     = {191--201},
  volume    = {80},
  doi       = {10.1007/s00248-019-01477-6},
}

@Article{SVH+19,
  author    = {Sollai, Martina and Villanueva, Laura and Hopmans, Ellen C. and Reichart, Gert-Jan and Sinninghe Damst\'e, Jaap S.},
  journal   = {Geobiology},
  title     = {A combined lipidomic and 16{S} r{RNA} gene amplicon sequencing approach reveals archaeal sources of intact polar lipids in the stratified {B}lack {S}ea water column},
  year      = {2019},
  number    = {1},
  pages     = {91--109},
  volume    = {17},
  doi       = {10.1111/gbi.12316},
}

@Article{HXZ+20,
  author    = {He, Peiqing and Xie, Linping and Zhang, Xuelei and Li, Jiang and Lin, Xuezheng and Pu, Xinming and Yuan, Chao and Tian, Ziwen and Li, Jie},
  journal   = {Scientific Reports},
  title     = {Microbial diversity and metabolic potential in the stratified {S}ansha {Y}ongle {B}lue {H}ole in the {S}outh {C}hina {S}ea},
  year      = {2020},
  number    = {1},
  pages     = {5949},
  volume    = {10},
  doi       = {10.1038/s41598-020-62411-2},
}

@Article{XWP+19,
  author    = {Linping Xie and Baodong Wang and Xinming Pu and Ming Xin and Peiqing He and Chengxuan Li and Qinsheng Wei and Xuelei Zhang and Tiegang Li},
  journal   = {Science of The Total Environment},
  title     = {Hydrochemical properties and chemocline of the {S}ansha {Y}ongle {B}lue {H}ole in the {S}outh {C}hina {S}ea},
  year      = {2019},
  pages     = {1281--1292},
  volume    = {649},
  doi       = {10.1016/j.scitotenv.2018.08.333},
}

@Article{ZML+17,
  author    = {Zhou, Zhichao and Meng, Han and Liu, Yang and Gu, Ji-Dong and Li, Meng},
  journal   = {Frontiers in Microbiology},
  title     = {Stratified bacterial and archaeal community in mangrove and intertidal wetland mudflats revealed by high throughput 16{S} r{RNA} gene sequencing},
  year      = {2017},
  pages     = {2148},
  volume    = {8},
  doi       = {10.3389/fmicb.2017.02148},
}

@Article{BCA+21,
  author    = {Baricz, Andreea and Chiriac, Cecilia Maria and Andrei, Adrian-{\c S}tefan and Bulzu, Paul-Adrian and Levei, Erika Andrea and Cadar, Oana and Battes, Karina Paula and C{\^i}mpean, Mirela and {\c S}enil{\u a}, Marin and Cristea, Adorj{\'a}n and Muntean, Vasile and Alexe, Mircea and Coman, Cristian and Szekeres, Edina Kriszta and Sicora, Cosmin Ionel and Ionescu, Artur and Blain, David and O'Neill, William Kenneth and Edwards, Jessica and Hallsworth, John Edward and Banciu, Horia Leonard},
  journal   = {Environmental Microbiology},
  title     = {Spatio-temporal insights into microbiology of the freshwater-to-hypersaline, oxic-hypoxic-euxinic waters of {U}rsu {L}ake},
  year      = {2021},
  number    = {7},
  pages     = {3523--3540},
  volume    = {23},
  doi       = {10.1111/1462-2920.14909},
}

@Article{MLL+19,
  author    = {Delong Meng and Juan Li and Tianbo Liu and Yongjun Liu and Mingli Yan and Jin Hu and Xiaoqi Li and Xueduan Liu and Yili Liang and Hongwei Liu and Huaqun Yin},
  journal   = {Journal of Environmental Sciences},
  title     = {Effects of redox potential on soil cadmium solubility: Insight into microbial community},
  year      = {2019},
  pages     = {224--232},
  volume    = {75},
  doi       = {10.1016/j.jes.2018.03.032},
}

@Article{LSS11,
  author    = {Leinonen, Rasko and Sugawara, Hideaki and Shumway, Martin},
  journal   = {Nucleic Acids Research},
  title     = {The {S}equence {R}ead {A}rchive},
  year      = {2011},
  number    = {suppl\_1},
  pages     = {D19--D21},
  volume    = {39},
  doi       = {10.1093/nar/gkq1019},
}

@Article{RSJ+21,
  author    = {Rissanen, Antti J. and Saarela, Taija and Jäntti, Helena and Buck, Moritz and Peura, Sari and Aalto, Sanni L. and Ojala, Anne and Pumpanen, Jukka and Tiirola, Marja and Elvert, Marcus and Nykänen, Hannu},
  journal   = {FEMS Microbiology Ecology},
  title     = {Vertical stratification patterns of methanotrophs and their genetic controllers in water columns of oxygen-stratified boreal lakes},
  year      = {2021},
  number    = {2},
  pages     = {fiaa252},
  volume    = {97},
  doi       = {10.1093/femsec/fiaa252},
}

@Article{NTB+21,
  author    = {Nothaft, Daniel and Templeton, Alexis S. and Boyd, Eric and Matter, Juerg and Stute, Martin and Paukert Vankeuren, Amelia N.},
  journal   = {Earth and Space Science Open Archive},
  title     = {Aqueous geochemical and microbial variation across discrete depth intervals in a peridotite aquifer assessed using a packer system in the {S}amail {O}phiolite, {O}man},
  year      = {2021},
  doi       = {10.1002/essoar.10506402.2},
}

@Article{RMB+17,
  author    = {Rempfert, Kaitlin R. and Miller, Hannah M. and Bompard, Nicolas and Nothaft, Daniel and Matter, Juerg M. and Kelemen, Peter and Fierer, Noah and Templeton, Alexis S.},
  journal   = {Frontiers in Microbiology},
  title     = {Geological and geochemical controls on subsurface microbial life in the {S}amail {O}phiolite, {O}man},
  year      = {2017},
  pages     = {56},
  volume    = {8},
  doi       = {10.3389/fmicb.2017.00056},
}

@Article{SBP+20,
  author    = {Sabuda, Mary C. and Brazelton, William J. and Putman, Lindsay I. and McCollom, Tom M. and Hoehler, Tori M. and Kubo, Michael D. Y. and Cardace, Dawn and Schrenk, Matthew O.},
  journal   = {Environmental Microbiology},
  title     = {A dynamic microbial sulfur cycle in a serpentinizing continental ophiolite},
  year      = {2020},
  number    = {6},
  pages     = {2329--2345},
  volume    = {22},
  doi       = {10.1111/1462-2920.15006},
}

@Article{CTS+17,
  author    = {Crespo-Medina, Melitza and Twing, Katrina I. and Sánchez-Murillo, Ricardo and Brazelton, William J. and McCollom, Thomas M. and Schrenk, Matthew O.},
  journal   = {Frontiers in Microbiology},
  title     = {Methane dynamics in a tropical serpentinizing environment: The {S}anta {E}lena {O}phiolite, {C}osta {R}ica},
  year      = {2017},
  pages     = {916},
  volume    = {8},
  doi       = {10.3389/fmicb.2017.00916},
}

@Article{MWY+21,
  author    = {Ma, Li and Wu, Geng and Yang, Jian and Huang, Liuqin and Phurbu, Dorji and Li, Wen-Jun and Jiang, Hongchen},
  journal   = {Frontiers in Microbiology},
  title     = {Distribution of hydrogen-producing bacteria in {T}ibetan hot springs, {C}hina},
  year      = {2021},
  pages     = {1789},
  volume    = {12},
  doi       = {10.3389/fmicb.2021.569020},
}

@Article{HSF+19,
  author    = {von Hoyningen-Huene, Avril Jean Elisabeth and Schneider, Dominik and Fussmann, Dario and Reimer, Andreas and Arp, Gernot and Daniel, Rolf},
  journal   = {Scientific Data},
  title     = {Bacterial succession along a sediment porewater gradient at {L}ake {N}eusiedl in {A}ustria},
  year      = {2019},
  number    = {1},
  pages     = {163},
  volume    = {6},
  doi       = {10.1038/s41597-019-0172-9},
}

@Article{SCM+18,
  author    = {Sánchez-Soto Jiménez, Ma. Fernanda and Cerqueda-García, Daniel and Montero-Muñoz, Jorge L. and Aguirre-Macedo, Ma. Leopoldina and García-Maldonado, José Q. and Souza, Valeria},
  journal   = {PeerJ},
  title     = {Assessment of the bacterial community structure in shallow and deep sediments of the {P}erdido {F}old {B}elt region in the {G}ulf of {M}exico},
  year      = {2018},
  pages     = {e5583},
  volume    = {6},
  doi       = {10.7717/peerj.5583},
}

@Article{BOEM21,
  author    = {Block, Kaleigh R. and O'Brien, Joy M. and Edwards, William J. and Marnocha, Cassandra L.},
  journal   = {MicrobiologyOpen},
  title     = {Vertical structure of the bacterial diversity in meromictic {F}ayetteville {G}reen {L}ake},
  year      = {2021},
  number    = {4},
  pages     = {e1228},
  volume    = {10},
  doi       = {10.1002/mbo3.1228},
}

@Article{HDZ+19,
  author    = {Maozhen Han and Melissa Dsouza and Chunyu Zhou and Hongjun Li and Junqian Zhang and Chaoyun Chen and Qi Yao and Chaofang Zhong and Hao Zhou and Jack A. Gilbert and Zhi Wang and Kang Ning},
  journal   = {Genomics, Proteomics \& Bioinformatics},
  title     = {Agricultural risk factors influence microbial ecology in {H}onghu {L}ake},
  year      = {2019},
  number    = {1},
  pages     = {76--90},
  volume    = {17},
  doi       = {10.1016/j.gpb.2018.04.008},
}

@Article{YHK+20,
  author    = {Lijuan Yan and Martina Herrmann and Bernd Kampe and Robert Lehmann and Kai Uwe Totsche and Kirsten Küsel},
  journal   = {Water Research},
  title     = {Environmental selection shapes the formation of near-surface groundwater microbiomes},
  year      = {2020},
  pages     = {115341},
  volume    = {170},
  doi       = {10.1016/j.watres.2019.115341},
}

@Article{ZHZ+19,
  author    = {Kun Zhang and Zheng Hu and Feifan Zeng and Xingjian Yang and Jinjin Wang and Ran Jing and Huanni Zhang and Yongtao Li and Zhen Zhang},
  journal   = {Journal of Environmental Management},
  title     = {Biodegradation of petroleum hydrocarbons and changes in microbial community structure in sediment under nitrate-, ferric-, sulfate-reducing and methanogenic conditions},
  year      = {2019},
  pages     = {109425},
  volume    = {249},
  doi       = {10.1016/j.jenvman.2019.109425},
}

@Article{BMJ+19,
  author    = {Belilla, Jodie and Moreira, David and Jardillier, Ludwig and Reboul, Guillaume and Benzerara, Karim and López-García, José M. and Bertolino, Paola and López-Archilla, Ana I. and López-García, Purificación},
  journal   = {Nature Ecology \& Evolution},
  title     = {Hyperdiverse archaea near life limits at the polyextreme geothermal {D}allol area},
  year      = {2019},
  number    = {11},
  pages     = {1552--1561},
  volume    = {3},
  doi       = {10.1038/s41559-019-1005-0},
}

@Article{CNA+20,
  author    = {Chopyk, Jessica and Nasko, Daniel J. and Allard, Sarah and Bui, Anthony and Pop, Mihai and Mongodin, Emmanuel F. and Sapkota, Amy R.},
  journal   = {Environmental Microbiome},
  title     = {Seasonal dynamics in taxonomy and function within bacterial and viral metagenomic assemblages recovered from a freshwater agricultural pond},
  year      = {2020},
  number    = {1},
  pages     = {18},
  volume    = {15},
  doi       = {10.1186/s40793-020-00365-8},
}

@Article{SRM+19,
  author    = {Sonthiphand, Prinpida and Ruangroengkulrith, Siwat and Mhuantong, Wuttichai and Charoensawan, Varodom and Chotpantarat, Srilert and Boonkaewwan, Satika},
  journal   = {Environmental Science and Pollution Research},
  title     = {Metagenomic insights into microbial diversity in a groundwater basin impacted by a variety of anthropogenic activities},
  year      = {2019},
  number    = {26},
  pages     = {26765--26781},
  volume    = {26},
  doi       = {10.1007/s11356-019-05905-5},
}

@Article{HLZ+18,
  author    = {Anyi Hu and Shuang Li and Lanping Zhang and Hongjie Wang and Jun Yang and Zhuanxi Luo and Azhar Rashid and Shaoqing Chen and Weixiong Huang and Chang-Ping Yu},
  journal   = {Environmental Pollution},
  title     = {Prokaryotic footprints in urban water ecosystems: A case study of urban landscape ponds in a coastal city, {C}hina},
  year      = {2018},
  pages     = {1729--1739},
  volume    = {242},
  doi       = {10.1016/j.envpol.2018.07.097},
}

@Article{XLD+20,
  author    = {Xu, Menglong and Liu, Yazi and Deng, Yan and Zhang, Siyuan and Hao, Xiaodong and Zhu, Ping and Zhou, Jieyi and Yin, Huaqun and Liang, Yili and Liu, Hongwei and Liu, Xueduan and Bai, Lianyang and Jiang, Luhua and Jiang, Huidan},
  journal   = {RSC Advances},
  title     = {Bioremediation of cadmium-contaminated paddy soil using an autotrophic and heterotrophic mixture},
  year      = {2020},
  number    = {44},
  pages     = {26090--26101},
  volume    = {10},
  doi       = {10.1039/D0RA03935G},
}

@Article{KSR+21,
  author    = {Kamran, Aysha and Sauter, Kathrin and Reimer, Andreas and Wacker, Theresa and Reitner, Joachim and Hoppert, Michael},
  journal   = {Microorganisms},
  title     = {Cyanobacterial mats in calcite-precipitating serpentinite-hosted alkaline springs of the {V}oltri {M}assif, {I}taly},
  year      = {2021},
  number    = {1},
  pages     = {62},
  volume    = {9},
  doi       = {10.3390/microorganisms9010062},
}

@Article{ZCZ+21,
  author    = {Zecchin, Sarah and Crognale, Simona and Zaccheo, Patrizia and Fazi, Stefano and Amalfitano, Stefano and Casentini, Barbara and Callegari, Matteo and Zanchi, Raffaella and Sacchi, Gian Attilio and Rossetti, Simona and Cavalca, Lucia},
  journal   = {Frontiers in Microbiology},
  title     = {Adaptation of microbial communities to environmental arsenic and selection of arsenite-oxidizing bacteria from contaminated groundwaters},
  year      = {2021},
  pages     = {364},
  volume    = {12},
  doi       = {10.3389/fmicb.2021.634025},
}

@Article{SKP+21,
  author    = {Prinpida Sonthiphand and Supeerapat Kraidech and Saowarod Polart and Srilert Chotpantarat and Kanthida Kusonmano and Pichahpuk Uthaipaisanwong and Chalida Rangsiwutisak and Ekawan Luepromchai},
  journal   = {Journal of Environmental Science and Health, Part A},
  title     = {Arsenic speciation, the abundance of arsenite-oxidizing bacteria and microbial community structures in groundwater, surface water, and soil from a gold mine},
  year      = {2021},
  number    = {7},
  pages     = {769--785},
  volume    = {56},
  doi       = {10.1080/10934529.2021.1927421},
}

@Article{ZZL+21,
  author    = {Lili Zhang and Cheng Zhang and Keting Lian and Dongfang Ke and Ting Xie and Chongxuan Liu},
  journal   = {Science of The Total Environment},
  title     = {River restoration changes distributions of antibiotics, antibiotic resistance genes, and microbial community},
  year      = {2021},
  pages     = {147873},
  volume    = {788},
  doi       = {10.1016/j.scitotenv.2021.147873},
}

@Article{PBU+20,
  author    = {Peltek, Sergey E. and Bryanskaya, Alla V. and Uvarova, Yuliya E. and Rozanov, Aleksey S. and Ivanisenko, Timofey V. and Ivanisenko, Vladimir A. and Lazareva, Elena V. and Saik, Olga V. and Efimov, Vadim M. and Zhmodik, Sergey M. and Taran, Oxana P. and Slynko, Nikolay M. and Shekhovtsov, Sergey V. and Parmon, Valentin N. and Dobretsov, Nikolay L. and Kolchanov, Nikolay A.},
  journal   = {BMC Microbiology},
  title     = {Young «oil site» of the {U}zon {C}aldera as a habitat for unique microbial life},
  year      = {2020},
  number    = {2},
  pages     = {349},
  volume    = {20},
  doi       = {10.1186/s12866-020-02012-1},
}

@Article{GWS+19,
  author    = {Liang Guo and Guangcai Wang and Yizhi Sheng and Zheming Shi and Xiaoyi Sun},
  journal   = {Science of The Total Environment},
  title     = {Groundwater microbial communities and their connection to hydrochemical environment in {G}olmud, {N}orthwest {C}hina},
  year      = {2019},
  pages     = {133848},
  volume    = {695},
  doi       = {10.1016/j.scitotenv.2019.133848},
}

@Article{KLY+20,
  author    = {Minkyung Kim and Jaebok Lee and Dongwoo Yang and Hye Yoon Park and Woojun Park},
  journal   = {Environmental Pollution},
  title     = {Seasonal dynamics of the bacterial communities associated with cyanobacterial blooms in the {H}an {R}iver},
  year      = {2020},
  pages     = {115198},
  volume    = {266},
  doi       = {10.1016/j.envpol.2020.115198},
}

@Article{SRM+21,
  author    = {Sonthiphand, Prinpida and Rattanaroongrot, Pasunun and Mek-yong, Kasarnchon and Kusonmano, Kanthida and Rangsiwutisak, Chalida and Uthaipaisanwong, Pichahpuk and Chotpantarat, Srilert and Termsaithong, Teerasit},
  journal   = {PeerJ},
  title     = {Microbial community structure in aquifers associated with arsenic: Analysis of 16{S} r{RNA} and arsenite oxidase genes},
  year      = {2021},
  pages     = {e10653},
  volume    = {9},
  doi       = {10.7717/peerj.10653},
}

@Article{JDP+20,
  author    = {Fadwa Jroundi and Michael Descostes and Cristina Povedano-Priego and Iván Sánchez-Castro and Vitthavat Suvannagan and Pierre Grizard and Mohamed L. Merroun},
  journal   = {Science of The Total Environment},
  title     = {Profiling native aquifer bacteria in a uranium roll-front deposit and their role in biogeochemical cycle dynamics: Insights regarding in situ recovery mining},
  year      = {2020},
  pages     = {137758},
  volume    = {721},
  doi       = {10.1016/j.scitotenv.2020.137758},
}

@Article{MLL+18,
  author    = {Mai, Yong-zhan and Lai, Zi-ni and Li, Xin-hui and Peng, Song-yao and Wang, Chao},
  journal   = {Marine Pollution Bulletin},
  title     = {Structural and functional shifts of bacterioplanktonic communities associated with spatiotemporal gradients in river outlets of the subtropical {P}earl {R}iver {E}stuary, {S}outh {C}hina},
  year      = {2018},
  pages     = {309--321},
  volume    = {136},
  doi       = {10.1016/j.marpolbul.2018.09.013},
}

@Article{SDH+19,
  author    = {Sang, Shilei and Dai, Heng and Hu, Bill X. and Hao, Yanru and Zhou, Tong and Zhang, Jin},
  journal   = {Water},
  title     = {The study of hydrogeochemical environments and microbial communities along a groundwater salinity gradient in the {P}earl {R}iver {D}elta, {C}hina},
  year      = {2019},
  number    = {4},
  pages     = {804},
  volume    = {11},
  doi       = {10.3390/w11040804},
}

@Article{BWD+19,
  author    = {Butler, Timothy M. and Wilhelm, Anna-Catharina and Dwyer, Amber C. and Webb, Paige N. and Baldwin, Andrew L. and Techtmann, Stephen M.},
  journal   = {Scientific Reports},
  title     = {Microbial community dynamics during lake ice freezing},
  year      = {2019},
  number    = {1},
  pages     = {6231},
  volume    = {9},
  doi       = {10.1038/s41598-019-42609-9},
}

@Article{LXH+20,
  author    = {Luo, Xia and Xiang, Xinyi and Huang, Guoyi and Song, Xiaorui and Wang, Peijia and Yang, Yuanhao and Fu, Kaidao and Che, Rongxiao},
  journal   = {Environmental Science and Pollution Research},
  title     = {Bacterial community structure upstream and downstream of cascade dams along the {L}ancang {R}iver in southwestern {C}hina},
  year      = {2020},
  number    = {34},
  pages     = {42933--42947},
  volume    = {27},
  doi       = {10.1007/s11356-020-10159-7},
}

@Article{MCC+20,
  author    = {Massello, Francisco L. and Chan, Chia Sing and Chan, Kok-Gan and Goh, Kian Mau and Donati, Edgardo and Urbieta, María Sofía},
  journal   = {Microorganisms},
  title     = {Meta-analysis of microbial communities in hot springs: Recurrent taxa and complex shaping factors beyond p{H} and temperature},
  year      = {2020},
  number    = {6},
  pages     = {906},
  volume    = {8},
  doi       = {10.3390/microorganisms8060906},
}

@Article{LMG+20,
  author    = {Lopez Bedogni, Germán and Massello, Francisco L. and Giaveno, Alejandra and Donati, Edgardo Rubén and Urbieta, María Sofía},
  journal   = {Microorganisms},
  title     = {A deeper look into the biodiversity of the extremely acidic {C}opahue {V}olcano-{R}ío {A}grio system in {N}euquén, {A}rgentina},
  year      = {2020},
  number    = {1},
  pages     = {58},
  volume    = {8},
  doi       = {10.3390/microorganisms8010058},
}

@Article{LLL+21,
  author    = {Xin Li and Xuefeng Li and Yueyue Li and Xiongze Dai and Qingzhuang Zhang and Mi Zhang and Zhuqing Zhang and Yu Tao and Wenchao Chen and Mingxing Zhang and Xiangyu Zhou and Sha Yang and Yanqing Ma and Mostafa Zhran and Xuexiao Zou},
  journal   = {Science of The Total Environment},
  title     = {Improved immobilization of soil cadmium by regulating soil characteristics and microbial community through reductive soil disinfestation},
  year      = {2021},
  pages     = {146222},
  volume    = {778},
  doi       = {10.1016/j.scitotenv.2021.146222},
}

@Article{WHL+21,
  author    = {Wang, Lixiao and Han, Maozhen and Li, Xi and Yu, Bingbing and Wang, Huading and Ginawi, Amjed and Ning, Kang and Yan, Yunjun},
  journal   = {Molecular Ecology},
  title     = {Mechanisms of niche-neutrality balancing can drive the assembling of microbial community},
  year      = {2021},
  number    = {6},
  pages     = {1492--1504},
  volume    = {30},
  doi       = {10.1111/mec.15825},
}

@Article{GWSS21,
  author    = {Guo, Liang and Wang, Guangcai and Sheng, Yizhi and Shi, Zheming},
  journal   = {Journal of Geophysical Research: Biogeosciences},
  title     = {Hydrogeochemical constraints shape hot spring microbial community compositions: Evidence from acidic, moderate-temperature springs and alkaline, high-temperature springs, southwestern {Y}unnan geothermal areas, {C}hina},
  year      = {2021},
  number    = {3},
  pages     = {e2020JG005868},
  volume    = {126},
  doi       = {10.1029/2020JG005868},
}

@Article{PCL+18,
  author    = {Power, Jean F. and Carere, Carlo R. and Lee, Charles K. and Wakerley, Georgia L. J. and Evans, David W. and Button, Mathew and White, Duncan and Climo, Melissa D. and Hinze, Annika M. and Morgan, Xochitl C. and McDonald, Ian R. and Cary, S. Craig and Stott, Matthew B.},
  journal   = {Nature Communications},
  title     = {Microbial biogeography of 925 geothermal springs in {N}ew {Z}ealand},
  year      = {2018},
  number    = {1},
  pages     = {2876},
  volume    = {9},
  doi       = {10.1038/s41467-018-05020-y},
}

@Article{RBW+14,
  author    = {Rundell, Ethan A. and Banta, Lois M. and Ward, Doyle V. and Watts, Corey D. and Birren, Bruce and Esteban, David J.},
  journal   = {PLOS One},
  title     = {16{S} r{RNA} gene survey of microbial communities in {W}inogradsky columns},
  year      = {2014},
  number    = {8},
  pages     = {e104134},
  volume    = {9},
  doi       = {10.1371/journal.pone.0104134},
}

@Article{DFYS19,
  author    = {Diez-Ercilla, Marta and Falagán, Carmen and Yusta, Iñaki and Sánchez-España, Javier},
  journal   = {Journal of Soils and Sediments},
  title     = {Metal mobility and mineral transformations driven by bacterial activity in acidic pit lake sediments: Evidence from column experiments and sequential extraction},
  year      = {2019},
  number    = {3},
  pages     = {1527--1542},
  volume    = {19},
  doi       = {10.1007/s11368-018-2112-2},
}

@Article{DTJ+20,
  author    = {Jun Dai and Zhu Tang and Nan Jiang and Peter M. Kopittke and Fang-Jie Zhao and Peng Wang},
  journal   = {Environmental Pollution},
  title     = {Increased arsenic mobilization in the rice rhizosphere is mediated by iron-reducing bacteria},
  year      = {2020},
  pages     = {114561},
  volume    = {263},
  doi       = {10.1016/j.envpol.2020.114561},
}

@Article{SBW+17,
  author    = {Yang Song and Yongrong Bian and Fang Wang and Anna Herzberger and Xinglun Yang and Chenggang Gu and Xin Jiang},
  journal   = {Chemosphere},
  title     = {Effects of biochar on dechlorination of hexachlorobenzene and the bacterial community in paddy soil},
  year      = {2017},
  pages     = {116--123},
  volume    = {186},
  doi       = {10.1016/j.chemosphere.2017.07.139},
}

@Article{WFB+21,
  author    = {Wyness, Adam J. and Fortune, Irene and Blight, Andrew J. and Browne, Patricia and Hartley, Morgan and Holden, Matthew and Paterson, David M.},
  journal   = {PLOS One},
  title     = {Ecosystem engineers drive differing microbial community composition in intertidal estuarine sediments},
  year      = {2021},
  number    = {2},
  pages     = {e0240952},
  volume    = {16},
  doi       = {10.1371/journal.pone.0240952},
}

@Article{BYB+17,
  author    = {Eric S. Boyd and Ri-Qing Yu and Tamar Barkay and Trinity L. Hamilton and Bonnie K. Baxter and David L. Naftz and Mark Marvin-DiPasquale},
  journal   = {Science of The Total Environment},
  title     = {Effect of salinity on mercury methylating benthic microbes and their activities in {G}reat {S}alt {L}ake, {U}tah},
  year      = {2017},
  pages     = {495--506},
  volume    = {581-582},
  doi       = {10.1016/j.scitotenv.2016.12.157},
}

@Article{KLM+16,
  author    = {Kocur, Chris M. D. and Lomheim, Line and Molenda, Olivia and Weber, Kela P. and Austrins, Leanne M. and Sleep, Brent E. and Boparai, Hardiljeet K. and Edwards, Elizabeth A. and O’Carroll, Denis M.},
  journal   = {Environmental Science \& Technology},
  title     = {Long-term field study of microbial community and dechlorinating activity following carboxymethyl cellulose-stabilized nanoscale zero-valent iron injection},
  year      = {2016},
  number    = {14},
  pages     = {7658--7670},
  volume    = {50},
  doi       = {10.1021/acs.est.6b01745},
}

@Article{LMBA21,
  author    = {Lanzén, Anders and Mendibil, Iñaki and Borja, {\'A}ngel and Alonso-Sáez, Laura},
  journal   = {Molecular Ecology},
  title     = {A microbial \emph{mandala} for environmental monitoring: Predicting multiple impacts on estuarine prokaryote communities of the {B}ay of {B}iscay},
  year      = {2021},
  number    = {13},
  pages     = {2969--2987},
  volume    = {30},
  doi       = {https://doi.org/10.1111/mec.15489},
}

@Article{ZDA+20,
  author    = {Zárate, Ana and Dorador, Cristina and Araya, Ruben and Guajardo, Mariela and Z. Florez, July and Icaza, Gonzalo and Cornejo, Diego and Valdés, Jorge},
  journal   = {PeerJ},
  title     = {Connectivity of bacterial assemblages along the {L}oa {R}iver in the {A}tacama {D}esert, {C}hile},
  year      = {2020},
  pages     = {e9927},
  volume    = {8},
  doi       = {10.7717/peerj.9927},
}

@Article{ZZZ+18,
  author    = {Zhu, Min and Zhang, Lujun and Zheng, Liwei and Zhuo, Ying and Xu, Jianming and He, Yan},
  journal   = {Frontiers in Microbiology},
  title     = {Typical soil redox processes in pentachlorophenol polluted soil following biochar addition},
  year      = {2018},
  pages     = {579},
  volume    = {9},
  doi       = {10.3389/fmicb.2018.00579},
}

@Article{BMOB18,
  author    = {Brigham, Brian A. and Montero, Angel D. and O'Mullan, Gregory D. and Bird, Jeffrey A.},
  journal   = {Soil Science Society of America Journal},
  title     = {Acetate additions stimulate {CO}\textsubscript{2} and {CH}\textsubscript{4} production from urban wetland soils},
  year      = {2018},
  number    = {5},
  pages     = {1147--1159},
  volume    = {82},
  doi       = {10.2136/sssaj2018.01.0034},
}

@Article{CWC+20,
  author    = {Shaochuang Chuang and Baozhan Wang and Kai Chen and Weibin Jia and Wenjing Qiao and Wanting Ling and Xiangyu Tang and Jiandong Jiang},
  journal   = {Science of The Total Environment},
  title     = {Microbial catabolism of lindane in distinct layers of acidic paddy soils combinedly affected by different water managements and bioremediation strategies},
  year      = {2020},
  pages     = {140992},
  volume    = {746},
  doi       = {10.1016/j.scitotenv.2020.140992},
}

@Article{JVW+20,
  author    = {Juhmani, Abdul-Salam and Vezzi, Alessandro and Wahsha, Mohammad and Buosi, Alessandro and Pascale, Fabio De and Schiavon, Riccardo and Sfriso, Adriano},
  journal   = {Microorganisms},
  title     = {Diversity and dynamics of seaweed associated microbial communities inhabiting the {L}agoon of {V}enice},
  year      = {2020},
  number    = {11},
  pages     = {1657},
  volume    = {8},
  doi       = {10.3390/microorganisms8111657},
}

@Article{GFE+16,
  author    = {Gallardo, Victor Ariel and Fonseca, Alexis and Espinoza, Carola and Ruiz-Tagle, Nathaly and Musleh, Selim},
  journal   = {Marine Biodiversity},
  title     = {Bacteria of the {H}umboldt sulfuretum comply with unifying macroecological principles},
  year      = {2016},
  number    = {2},
  pages     = {399--406},
  volume    = {46},
  doi       = {10.1007/s12526-015-0377-x},
}

@Article{LJC+20,
  author    = {Luan, Lu and Jiang, Yuji and Cheng, Menghua and Dini-Andreote, Francisco and Sui, Yueyu and Xu, Qinsong and Geisen, Stefan and Sun, Bo},
  journal   = {Nature Communications},
  title     = {Organism body size structures the soil microbial and nematode community assembly at a continental and global scale},
  year      = {2020},
  number    = {1},
  pages     = {6406},
  volume    = {11},
  doi       = {10.1038/s41467-020-20271-4},
}

@Article{ZZLL21,
  author    = {Lili Zhang and Cheng Zhang and Keting Lian and Chongxuan Liu},
  journal   = {Journal of Hazardous Materials},
  title     = {Effects of chronic exposure of antibiotics on microbial community structure and functions in hyporheic zone sediments},
  year      = {2021},
  pages     = {126141},
  volume    = {416},
  doi       = {10.1016/j.jhazmat.2021.126141},
}

@Article{DLS21,
  author    = {Dykes, Gretchen E. and Limmer, Matt A. and Seyfferth, Angelia L.},
  journal   = {Plant and Soil},
  title     = {Silicon-rich soil amendments impact microbial community composition and the composition of ars{M} bearing microbes},
  year      = {2021},
  doi       = {10.1007/s11104-021-05103-8},
}

@Article{FAV+21,
  author    = {Fazi, Stefano and Amalfitano, Stefano and Venturi, Stefania and Pacini, Nic and Vazquez, Eusebi and Olaka, Lydia A. and Tassi, Franco and Crognale, Simona and Herzsprung, Peter and Lechtenfeld, Oliver J. and Cabassi, Jacopo and Capecchiacci, Francesco and Rossetti, Simona and Yakimov, Michail M. and Vaselli, Orlando and Harper, David M. and Butturini, Andrea},
  journal   = {Communications Biology},
  title     = {High concentrations of dissolved biogenic methane associated with cyanobacterial blooms in {E}ast {A}frican lake surface water},
  year      = {2021},
  number    = {1},
  pages     = {845},
  volume    = {4},
  doi       = {10.1038/s42003-021-02365-x},
}

@Article{CLS+19,
  author    = {Wei Cai and Yi Li and Yun Shen and Chao Wang and Peifang Wang and Longfei Wang and Lihua Niu and Wenlong Zhang},
  journal   = {Journal of Environmental Management},
  title     = {Vertical distribution and assemblages of microbial communities and their potential effects on sulfur metabolism in a black-odor urban river},
  year      = {2019},
  pages     = {368--376},
  volume    = {235},
  doi       = {10.1016/j.jenvman.2019.01.078},
}

@Article{OFY+19,
  author    = {Omae, Kimiho and Fukuyama, Yuto and Yasuda, Hisato and Mise, Kenta and Yoshida, Takashi and Sako, Yoshihiko},
  journal   = {Archives of Microbiology},
  title     = {Diversity and distribution of thermophilic hydrogenogenic carboxydotrophs revealed by microbial community analysis in sediments from multiple hydrothermal environments in {J}apan},
  year      = {2019},
  number    = {7},
  pages     = {969--982},
  volume    = {201},
  doi       = {10.1007/s00203-019-01661-9},
}

@Article{MCS+21,
  author    = {Molina, Verónica and Cornejo-D’Ottone, Marcela and Soto, Eulogio H. and Quiroga, Eduardo and Alarcón, Guillermo and Silva, Daniela and Acuña, Carla and Silva, Nelson},
  journal   = {Water},
  title     = {Biogeochemical responses and seasonal dynamics of the benthic boundary layer microbial communities during the {E}l {N}iño 2015 in an eastern boundary upwelling system},
  year      = {2021},
  number    = {2},
  pages     = {180},
  volume    = {13},
  doi       = {10.3390/w13020180},
}

@Article{VMD+21,
  author    = {van Vliet, Daan M. and von Meijenfeldt, F.A. Bastiaan and Dutilh, Bas E. and Villanueva, Laura and Sinninghe Damsté, Jaap S. and Stams, Alfons J.M. and Sánchez-Andrea, Irene},
  journal   = {Environmental Microbiology},
  title     = {The bacterial sulfur cycle in expanding dysoxic and euxinic marine waters},
  year      = {2021},
  number    = {6},
  pages     = {2834--2857},
  volume    = {23},
  doi       = {10.1111/1462-2920.15265},
}

@Manual{KR21,
  title     = {{oce}: Analysis of Oceanographic Data},
  author    = {Dan Kelley and Clark Richards},
  note      = {{R} package version 1.4-0},
  year      = {2021},
  url       = {https://CRAN.R-project.org/package=oce},
}

@Article{PMM+20,
  author    = {Pronk, Geertje J. and Mellage, Adrian and Milojevic, Tatjana and Smeaton, Christina M. and Engel, Katja and Neufeld, Josh D. and Rezanezhad, Fereidoun and Van Cappellen, Philippe},
  journal   = {Vadose Zone Journal},
  title     = {Carbon turnover and microbial activity in an artificial soil under imposed cyclic drainage and imbibition},
  year      = {2020},
  number    = {1},
  pages     = {e20021},
  volume    = {19},
  doi       = {10.1002/vzj2.20021},
}

@Article{GRG+20,
  author    = {Christen L. Grettenberger and Rebecca L. Mccauley Rench and Danielle S. Gruen and Daniel B. Mills and Colin Carney and Jamie Brainard and Hiroshi Hamasaki and Ramses Ramirez and Yumiko Watanabe and Linda A. Amaral-Zettler and Hiroshi Ohmoto and Jennifer L. Macalady},
  journal   = {Geomicrobiology Journal},
  title     = {Microbial population structure in a stratified, acidic pit lake in the {I}berian {P}yrite {B}elt},
  year      = {2020},
  number    = {7},
  pages     = {623--634},
  volume    = {37},
  doi       = {10.1080/01490451.2020.1751748},
}

@Article{NLE+21,
  author    = {Ninio, Shira and Lupu, Achsa and Eckert, Werner and Ostrovsky, Ilia and Viner Mozzini, Yehudit and Sukenik, Assaf},
  journal   = {Freshwater Biology},
  title     = {Metalimnetic chlorophyll maxima in {L}ake {K}inneret - \emph{{C}hlorobium} revisited},
  year      = {2021},
  number    = {3},
  pages     = {468--480},
  volume    = {66},
  doi       = {10.1111/fwb.13653},
}

@Article{GSY+20,
  author    = {van der Graaf, Charlotte M. and Sánchez-España, Javier and Yusta, Iñaki and Ilin, Andrey and Shetty, Sudarshan A. and Bale, Nicole J. and Villanueva, Laura and Stams, Alfons J. M. and Sánchez-Andrea, Irene},
  journal   = {Microorganisms},
  title     = {Biosulfidogenesis mediates natural attenuation in acidic mine pit lakes},
  year      = {2020},
  number    = {9},
  pages     = {1275},
  volume    = {8},
  doi       = {10.3390/microorganisms8091275},
}

@Article{APV+20,
  author    = {Eduardo J. Aguilar-Rangel and Blanca L. Prado and María Soledad Vásquez-Murrieta and Paulina Estrada-de {los Santos} and Christina Siebe and Luisa I. Falcón and Jazmín Santillán and Rocío J. Alcántara-Hernández},
  journal   = {Journal of Contaminant Hydrology},
  title     = {Temporal analysis of the microbial communities in a nitrate-contaminated aquifer and the co-occurrence of anammox, n-damo and nitrous-oxide reducing bacteria},
  year      = {2020},
  pages     = {103657},
  volume    = {234},
  doi       = {10.1016/j.jconhyd.2020.103657},
}

@Article{LLC+19,
  author    = {Min Liu and Lemian Liu and Huihuang Chen and Zheng Yu and Jun R. Yang and Yuanyuan Xue and Bangqin Huang and Jun Yang},
  journal   = {Science of The Total Environment},
  title     = {Community dynamics of free-living and particle-attached bacteria following a reservoir \emph{{M}icrocystis} bloom},
  year      = {2019},
  pages     = {501--511},
  volume    = {660},
  doi       = {10.1016/j.scitotenv.2018.12.414},
}

@Article{WHC+19,
  author    = {Jiaqi Wang and Miaolian Hua and Chaoyang Cai and Jiajie Hu and Junren Wang and Hongrui Yang and Fang Ma and Haifeng Qian and Ping Zheng and Baolan Hu},
  journal   = {Applied and Environmental Microbiology},
  title     = {Spatial-temporal pattern of sulfate-dependent anaerobic methane oxidation in an intertidal zone of the {E}ast {C}hina {S}ea},
  year      = {2019},
  number    = {7},
  pages     = {e02638-18},
  volume    = {85},
  doi       = {10.1128/AEM.02638-18},
}

@TechReport{USGS10,
  author      = {{USGS}},
  institution = {U.S. Geological Survey},
  title       = {Great {L}akes and {W}atersheds {S}hapefiles},
  year        = {2010},
  note        = {Accessed on 2021-09-29},
  type        = {Science{B}ase {C}atalog},
  url         = {https://www.sciencebase.gov/catalog/item/530f8a0ee4b0e7e46bd300dd},
}

@Article{GZL21,
  author    = {Yu Gao and Wenlong Zhang and Yi Li},
  journal   = {Water Research},
  title     = {Microbial community coalescence: Does it matter in the {T}hree {G}orges {R}eservoir?},
  year      = {2021},
  pages     = {117638},
  volume    = {205},
  doi       = {10.1016/j.watres.2021.117638},
}

@Article{LZR+17,
  author    = {Li, Huabing and Zeng, Jin and Ren, Lijuan and Wang, Jianjun and Xing, Peng and Wu, Qinglong L.},
  journal   = {Limnology and Oceanography},
  title     = {Contrasting patterns of diversity of abundant and rare bacterioplankton in freshwater lakes along an elevation gradient},
  year      = {2017},
  number    = {4},
  pages     = {1570--1585},
  volume    = {62},
  doi       = {10.1002/lno.10518},
}

@Article{WHLH21,
  author    = {Jiapeng Wu and Yiguo Hong and Xiaohan Liu and Yaohao Hu},
  journal   = {Environmental Pollution},
  title     = {Variations in nitrogen removal rates and microbial communities over sediment depth in {D}aya {B}ay, {C}hina},
  year      = {2021},
  pages     = {117267},
  volume    = {286},
  doi       = {10.1016/j.envpol.2021.117267},
}

@Article{YHK+19,
  author    = {Yamamoto, Kyosuke and Hackley, Keith C. and Kelly, Walton R. and Panno, Samuel V. and Sekiguchi, Yuji and Sanford, Robert A. and Liu, Wen-Tso and Kamagata, Yoichi and Tamaki, Hideyuki},
  journal   = {Scientific Reports},
  title     = {Diversity and geochemical community assembly processes of the living rare biosphere in a sand-and-gravel aquifer ecosystem in the {M}idwestern {U}nited {S}tates},
  year      = {2019},
  number    = {1},
  pages     = {13484},
  volume    = {9},
  doi       = {10.1038/s41598-019-49996-z},
}
